# Non-invasive modulation of the human dorsal anterior cingulate attenuates acute pain perception and homeostatic cardiovascular responses

**DOI:** 10.1101/2023.06.30.547251

**Authors:** Andrew Strohman, Brighton Payne, Alexander In, Katelyn Stebbins, Wynn Legon

## Abstract

Homeostasis is the process of maintaining physiologic balance in the body that is critical for maintaining health and is dysfunctional in several disorders like chronic pain. The dorsal anterior cingulate cortex (dACC) is a critical brain area for homeostatic cardiovascular responses and pain processing, making it a promising non-invasive therapeutic target. We leverage the high spatial resolution and deep focal lengths of low-intensity focused ultrasound (LIFU) to non-invasively modulate the dACC for an effect on behavioral and cardiac autonomic responses using a transient heat pain stimulus. N = 16 healthy human volunteers (6M/10F) received transient contact heat pain during either LIFU to the dACC or Sham stimulation. Continuous electroencephalogram (EEG), electrocardiogram (ECG), and electrodermal response (EDR) were recorded. Outcome measures included perceived pain ratings, homeostatic measures including heart-rate variability, blood pressure, EDR response as well as the amplitude of the contact heat-evoked potential (CHEP).

LIFU reduced pain ratings by 1.08 ± 0.21 points relative to Sham. LIFU increased heart rate variability indexed by the standard deviation of normal sinus beats (SDNN), low frequency (LF) power, and the low-frequency/high-frequency (LF/HF) ratio. There were no effects on blood pressure or EDR. LIFU resulted in a 25.1% reduction in the N1-P1 CHEP amplitude driven primarily by effects on the P1 amplitude. Our results demonstrate LIFU to the dACC reduces perceived pain and alters homeostatic cardiovascular responses to a transient heat pain stimulus. These results have implications for the causal understanding of human pain and autonomic processing in the dACC and the potential for future therapeutics for pain relief and homeostatic modulation.

**SIGNIFICANCE STATEMENT:** New lines of inquiry now demonstrate cardiac homeostatic signals like heart rate variability (HRV) are aberrant in mental health disorders, addiction, and chronic pain and may contribute to their underlying etiology. The dorsal anterior cingulate cortex (dACC) is a key homeostatic center with direct influences on cardiovascular autonomic function, but its depth precludes direct access without invasive surgery. For the first time in humans, we demonstrate low-intensity focused ultrasound (LIFU) can non-invasively and selectively modulate the dACC to reduce acute pain perception and homeostatic cardiovascular responses as well as pain processing signals. This work helps establish a causal role of the dACC in pain perception and homeostatic signaling with potential future clinical applications in chronic pain and neuropsychological populations.

## INTRODUCTION

Homeostasis consists of autonomic, hormonal, and behavioral mechanisms that maintain physiologic balance in the body, and the sense of these physiologic conditions is termed interoception (Craig, 2002, 2009). The Lamina I spinothalamic system is a key homeostatic pathway, transmitting information on the physiologic state of all tissues in the body through the thalamus to cortical regions including the insula and cingulate (Craig, 2002; Critchley, 2005). Nociceptive signals selectively transmit along this pathway and integrates information with the sympathetic cell columns and homeostatic regions like the ventrolateral medulla, parabrachial nucleus, and periaqueductal grey, leading to the experience of pain and reconceptualizing pain as a homeostatic signal (Craig, 2003). Evidence implicates dysfunctional homeostatic/interoceptive regulation across neuropsychological and chronic pain conditions (Khalsa et al., 2018; Bonaz et al., 2021).

The dorsal anterior cingulate cortex (dACC) receives direct spinothalamocortical projections and is a critical region for pain and homeostatic processing, making it a promising non-invasive therapeutic target (Beissner et al., 2013; Wager et al., 2013; Vogt, 2016; Seeley, 2019). Increased activity in the dACC is associated with pain and emotional processing (Shackman et al., 2011) and it encodes both pain intensity and affect (Wager et al., 2013; Xiao and Zhang, 2018). Overactivity in the dACC appears to be involved in the onset and maintenance of chronic pain conditions (Bliss et al., 2016; Vanneste and De Ridder, 2021).

### dACC and Autonomic Function

Pain is a potent autonomic challenge, and the autonomic nervous system functions to maintain homeostasis (Quadt et al., 2018). The dACC integrates cognitive processes, pain, and autonomic control (Shackman et al., 2011; Lee et al., 2020), and direct electrical stimulation generates autonomic and behavioral responses like tachycardia and heat sensation (Xue et al., 2023). Activity in the dACC is implicated in low-frequency (LF) power (Critchley et al., 2003; Rebollo et al., 2018), a measure of heart rate variability that indexes sympathetic and parasympathetic systems and baroreflex sensitivity (Goldstein et al., 2011; Reyes Del Paso et al., 2013). Baroreflexes play a role in maintaining homeostasis and pain suppression (Dworkin et al., 1994; Chung et al., 2008; Suarez-Roca et al., 2019) and other research directly implicates the dACC in baroreflex activity (Gianaros et al., 2012; Ginty et al., 2013). The dACC is thus a critical hub for autonomic function and pain processing, suggesting homeostasis and perceived pain are inextricably linked in chronic pain populations (Medford and Critchley, 2010; Seifert et al., 2013).

### Non-Invasive dACC Modulation and Low-intensity Focused Ultrasound

Non-invasive neuromodulation techniques like transcranial magnetic stimulation (TMS) and transcranial electrical stimulation (TES) have previously attempted to target the dACC with equivocal results (Galhardoni et al., 2019; Khan et al., 2020), likely because these technologies do not confer the spatial resolution or depth penetration to safely reach targets deeper than approximately 3cm in the brain (Deng et al., 2013a; O’Connell et al., 2018; Gomez-Tames et al., 2020). Low-intensity focused ultrasound (LIFU) is a neuromodulatory approach that uses mechanical energy to non-destructively and reversibly modulate neuronal activity with millimeter-scale spatial resolution and adjustable depth of focus suitable for targeting deeper brain targets with high spatial precision (Legon et al., 2014, 2018a; Blackmore et al., 2019; Darmani et al., 2022). LIFU has been effectively and safely used for cortical and sub-cortical neuromodulation in humans, making it the only available non-invasive tool capable of targeting the dACC (Legon et al., 2014, 2018a, 2018b; Cain et al., 2021; Nakajima et al., 2022).

It is the purpose of this paper to examine how modulation of the dACC with LIFU affects subjective report of pain, homeostatic cardiovascular responses and pain processing signals in response to transient painful contact heat. We hypothesize LIFU will confer an inhibitory effect (Legon et al., 2014, 2023) to reduce subjective pain perception, attenuate the contact heat evoked potential (CHEP) and cardiovascular response to a painful stimulus.

## MATERIALS AND METHODS

### Participants

A total of 16 healthy participants (6M/10F, aged 28 ± 5.46 years) met inclusion criteria and were enrolled in the study. Inclusion criteria were: males and females aged 18 – 65. Exclusion criteria were in accordance with contraindications to non-invasive neuromodulation as outlined by Rossi et al., 2009 (Rossi et al., 2009) for transcranial magnetic stimulation in addition to contraindications to MRI and CT; pregnancy; an active medical disorder or treatment with potential CNS effects or a history of neurologic disorder (e.g. Multiple Sclerosis, Parkinson’s, Epilepsy, etc.); a history of head injury resulting in loss of consciousness for >10 minutes and/or a history of alcohol or drug dependence. All procedures were reviewed and approved by the Virginia Tech Institutional Review Board.

### Overall Study Design

Data was collected in three sessions on three separate days with a minimum of 72 hours between sessions. Session 1 was an imaging visit consisting of an anatomical computerized tomography (CT) and structural magnetic resonance imaging (MRI). Sessions 2 and 3 were formal testing days where participants were counterbalanced for Sham or real LIFU to the dACC.

### Imaging Acquisition

MRI data were acquired on a Siemens 3T Prisma scanner (Siemens Medical Solutions, Erlangen, Germany) at the Fralin Biomedical Research Institute’s Human Neuroimaging Laboratory. Anatomical scans were acquired using a T1-weighted MPRAGE sequence with a TR = 1400 ms, TI = 600 ms, TE = 2.66 ms, flip angle = 12°, voxel size = 0.5×0.5×1.0 mm, FoV read = 245 mm, FoV phase of 87.5%, 192 slices, ascending acquisition. CT scans were collected with a Kernel = Hr60 in the bone window, FoV = 250 mm, kilovolts (kV) = 120, rotation time = 1 second, delay = 2 seconds, pitch = 0.55, caudocranial image acquisition order, 1.0 mm image increments for a total of 121 images and total scan time of 13.14 seconds.

### Study Procedures

#### Overview of testing procedures and timeline

Prior to formal testing for sessions 2 and 3, participants first completed a baseline ROS questionnaire that asks about the presence and severity of symptoms (sleepiness, headache etc.) as previously used in our human LIFU studies (Legon et al., 2020) (see Appendix for Questionnaire). They were then seated in a comfortable chair with arm and neck support and connected to continuous electroencephalogram (EEG), electrocardiogram (ECG), and electrodermal response (EDR) monitoring (see below for details). Pain thresholding (see below), baseline blood pressure and a 5 minute baseline EEG, ECG and EDR were then performed. After the 5-minute baseline, formal contact heat pain testing was performed which lasted about 10 minutes. After pain testing, participants rested for 5 minutes. Continuous EEG, ECG, and EDR data was collected throughout all time windows. After 30 minutes, participants completed another ROS questionnaire and an Auditory Masking Questionnaire (AMQ). Details of each procedure are below. The study timeline is depicted in **Figure 1**.

**Figure 1.**
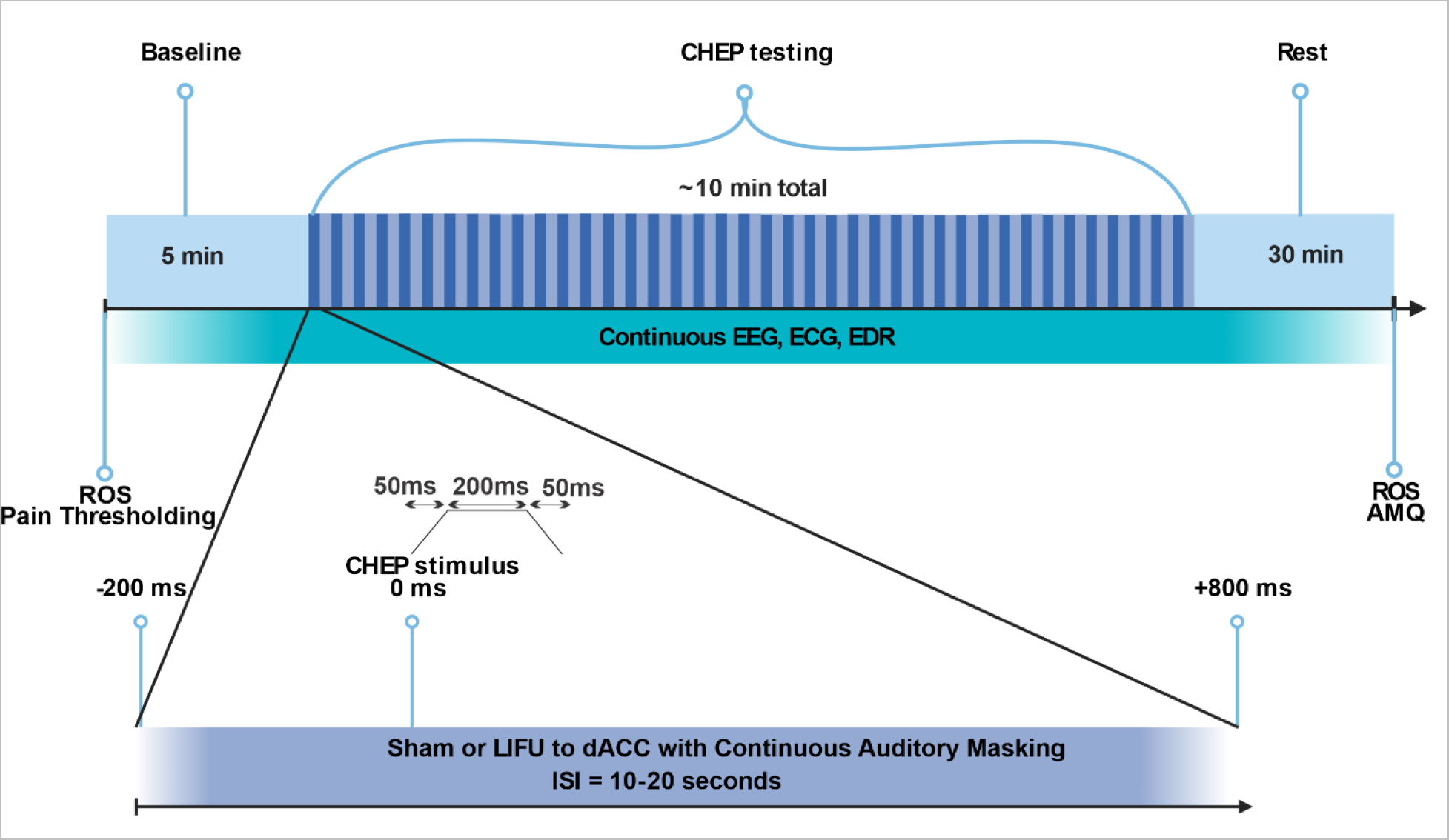
Timeline of pain testing. Data collection was divided into 3 time windows: A 5-minute baseline, 10-minute CHEP testing, and a 30-minute rest/waiting period. Prior to pain thresholding, a report of symptoms (ROS) questionnaire was administered followed by a 5 minute baseline period. CHEP testing consisted of 40 CHEP stimuli (300ms duration (50ms ramp, 200ms plateau, 50ms return) delivered to the dorsum of the right hand with a random inter-stimulus interval (ISI) of 10-20 seconds. One second of low-intensity focused ultrasound (LIFU) was time-locked to the CHEP stimulus, beginning 200 milliseconds (ms) prior to the CHEP stimulus (0 ms) and ending 800ms after. Continuous electroencephalogram (EEG), electrocardiogram (ECG), and electrodermal response (EDR) were recorded throughout all time windows. After the 30 minute rest/wait period participants completed the auditory masking questionnaire (AMQ) and the second ROS.

#### Data Acquisition

*Electroencephalogram (EEG).* Surface EEG was collected from channels CZ and PZ. The vertex (CZ) has previously been shown as the best site for recording the CHEP (Granovsky et al., 2005; Chen et al., 2006). Data were continuously acquired at a 1 kHz sampling rate using a DC amplifier (GES 400, Magstim EGI, OR, USA) and Net Station™ 5.4 EEG software with two 10-mm silver-over-silver chloride cup electrodes, referenced to the bilateral mastoid processes. The scalp for each site was first prepared with a mild abrasive gel (Nuprep; Weaver and Company, Aurora, CO) and rubbing alcohol. Cup electrodes were filled with a conductive paste (Ten20 Conductive; Weaver and Company, Aurora, CO) and held in place with medical tape. Electrode impedances were verified (<50 kΩ) before recording. Data was stored on a PC for offline data analysis.

*Electrocardiogram (ECG).* Two latex-free ECG electrodes (MedGel™, MDSM611903) were attached symmetrically to the anterior surface of the bilateral forearms immediately distal to the antecubital fossa. ECG data was continuously collected and sampled at 1 kHz using the same 64-channel EEG recording system with integrated grounding (GES 400, Magstim EGI) with a physiologic data acquisition box (Physio16, Magstim, EGI) and Net Station™ 5.4 EEG software and stored on a PC for offline data analysis.

*Electrodermal Response (EDR).* EDR was collected using the Consensys™ GSR Development Kit (Shimmer, Cambridge, MA, USA). It was placed on the right wrist with a photoplethysmogram sensor on the proximal 4^th^ digit and skin conductance electrodes on the proximal 2^nd^ and 3^rd^ digits. EDR data was continuously collected and sampled at 128 Hz using the Consensys software and stored on a PC for offline data analysis.

*Blood Pressure (BP).* Systolic and diastolic BP were collected using the QardioArm™ wireless portable blood pressure monitor placed on the upper left arm. BP measurements were taken twice, once at the beginning of each session prior to any intervention or task (pre-session BP), and once at the end of each session after all session data was collected (post-session BP).

#### Contact Heat Pain Stimulus

The contact heat pain stimulus was delivered using the TCS (thermal cutaneous stimulator) system (QST.lab, Strasbourg, FR) and a contact 3×3.2×2.4 mm peltier device (T-03). The peltier device used for contact heat-evoked potential (CHEP) stimulus delivery is capable of delivering rapid ramping heat stimuli that actives peripheral Aδ-fibres but not C-fibers (Truini et al., 2007, Anon, 2018) and is a validated way of studying nociception and pain perception in both healthy (Chen et al., 2001, 2006; Granovsky et al., 2016) and chronic pain populations (Lenoir et al., 2020). This QST device is a validated tool to elicit the CHEP (De Schoenmacker et al., 2022).

#### Pain Thresholding

Prior to formal pain thresholding, participants were familiarized with the QST probe and the heat stimulus. Formal thresholding consisted of delivering 3 consecutive stimuli (as above starting at 55 °C) at a fixed ISI of 5 seconds. Participants were required to rate the perceived pain of each stimulus on a 0-9 numerical rating scale where 0 = no pain; 1-3 = mild pain; 4-6 = moderate pain; 7-9 = severe pain. The average of the three stimuli was used as their threshold. The temperataure was raised in 1°C increments until a 5/9 average rating was achieved. Thresholding was performed this way because pilot data indicated that thresholding using a continuous graded stimulus did not correlate with thresholds using the discrete 300 msec stimulus. We found using the actual CHEP stimulus as the thresholding stimulus produced more consistent pain ratings during formal testing.

#### CHEP Testing

Six points were first identified in a 3×2 cm grid (1 cm spacing) on the dorsum of the right hand to guide the location of the heat probe on the hand. Each CHEP stimulation consisted of stimulus duration of 0.3 seconds (50ms ramp up, 200ms plateau, 50ms ramp down) at a ramping speed of 300°C/s. In between each stimulus there was a random inter-stimulus interval (ISI) of 10-20 seconds. The probe was moved in a clockwise direction to the next spot on the 3×2 grid following each stimulus in order to minimize repeated stimuli at a single site and prevent habituation or sensitization to the CHEP stimulus at any one hand site. A total of 40 stimuli were delivered. Participants were required to rate their perceived pain on the 0-9 numerical rating scale using a numerical keypad in their left hand. There was no time restriction placed on the response. Pain ratings were recorded using a custom Matlab script using MATLAB R2022a (The MathWorks, Inc., Natick, MA).

#### LIFU Transducer and Waveform

We used a single-element focused ultrasound transducer (Sonic Concepts™, model H-104) with a center frequency of 0.5 MHz, 64 mm aperture, 52 mm focal length from the exit plane and an f# of 0.81. Transcranial ultrasonic neuromodulation waveforms were generated using a 30-MHz dual channel function generator (4078B; BK Precision Instruments). Channel 1 was a 5Vp-p square wave burst of 1kHz (N = 1000) with a 36% duty cycle used to gate channel 2 that was a 500 kHz sine wave. This resulted in a 1 second waveform of 1000 pulses of 360 microseconds (μs). The output of channel 2 was sent through a 100-W linear RF amplifier (E&I 2100L; Electronics & Innovation) before being sent to the transducer through its corresponding matching network. LIFU delivery was time-locked to the CHEP stimulus, beginning at −200 ms (prior to stimulus) and ending at +800 ms after heat stimulus delivery. This time-locked duration was chosen due to sex and age variability in CHEP latency (Granovsky et al., 2016), ensuring the CHEP was generated within the LIFU delivery.

#### Empirical Acoustic Field Mapping

The acoustic intensity profile of the ultrasound waveform was measured in an acoustic test tank filled with deionized, degassed, and filtered water (Precision Acoustics Ltd., Dorchester, Dorset, UK). A calibrated hydrophone (HNR-0500, Onda Corp., Sunnyvale, CA, USA) mounted on a motorized stage was used to measure the pressure from the ultrasound transducer in the acoustic test tank. The ultrasound transducer was positioned in the tank using a custom set-up and levelled to ensure the hydrophone was perpendicular to the surface of the transducer. XY and YZ planar scans were performed at an isotropic 0.25 × 0.25 mm resolution. A voltage sweep from input voltages of 20-250 mVpp in 10mVpp increments were also performed to determine the necessary input voltage to obtain the desired extracranial pressure.

#### LIFU Targeting

The Montreal Neurologic Institute (MNI) XYZ coordinates (0,18,30) were chosen based upon Neurosynth reverse inference software for the term “pain” as well as prior literature on the fMRI signature of heat pain (Yarkoni et al., 2011; Wager et al., 2013). This target was sufficiently anterior to allow placement of the EEG CZ electrode along with the transducer so that they did not overlap. LIFU targeting was aided using a neuronavigation system (BrainSight™, Rogue Research, Montreal, QUE, CAN). Individual subject T1-weighted structural MRIs were registered in the Brainsight™ neuronavigation software. Participant dACC targets were transformed from standard MNI space to individual space using the software’s native transformation so that the coordinates applied to each subject’s individual anatomy. In one subject (male subject 4 in **Table S1**) the native transformation placed the original MNI coordinate on the corpus callosum, so a secondary site (MNI: 0,26,26) was chosen as this allowed direct targeting of grey matter and is also a common activation site for thermal pain in Neurosynth (Yarkoni et al., 2011; Lieberman and Eisenberger, 2015; Wager et al., 2016). The LIFU transducer was coupled to the head by first parting the hair using standard ultrasound gel and then using an acoustically transparent gel puck (Strohman et al., 2023). Puck thickness was selected for each participant based on the distance from the scalp to the dACC target (**Table S1**) in order to individualize axial (depth) targeting between participants. Optimal positioning of the transducer was midline at the top of the head anterior to the vertex for all participants. The transducer was secured to the head and positioning verified online continuously throughout the experiment using the neuronavigation system’s native real-time tracking software.

#### Auditory Masking

LIFU is known to produce audible sounds that may confound the data, but these can be effectively masked (Braun et al., 2020; Liang et al., 2023). For auditory masking, participants were given headphones connected to a tablet with a multitone white noise generator. The sound options were a series of environmental sounds such as thunderstorms, traffic, wind chimes, etc. Participants were asked to select their sounds, including layering them on top of each other, and instructed to turn the volume up until they could not hear normal conversation. This was verified by querying them by standing behind them and verbally asking “can you hear me?” at a normal conversational level. A lack of response indicated an adequate volume level. The mask was constantly applied throughout the entire period of LIFU or Sham application. After each session, participants were queried on the effectiveness of auditory masking using the AMQ. **Auditory Masking Questionnaire.** The Auditory Masking Questionnaire (AMQ) was administered on both LIFU and Sham sessions after all testing was completed. It consisted of three questions to evaluate the success of auditory masking: “I could hear the LIFU stimulation,” “I could feel the LIFU stimulation,” and “I believe I experienced LIFU stimulation.” Each question was answered on a 7-point Likert scale where 1 = “Strongly Disagree,” 2 = “Disagree,” 3 = Slightly Disagree,”4 = “Unsure,” 5 = “Slightly Agree,” 6 = “Agree,” and 7= “Strongly Agree.”

#### Report of Symptoms (ROS) Questionnaire

The ROS asks about the presence of symptoms including severity (absent, mild, moderate, severe) and has been previously used in our human LIFU studies. All symptoms are scored on a 0-3 scale, with a 0 indicating absence of the symptom up to a 3 indicating a severe symptom. The ROS questionnaire collected before and after LIFU or Sham were summed across participants to provide a total count of absent, mild, moderate, and severe intensity reports for each symptom. Counts of symptoms after LIFU were adjusted by taking the pre-post LIFU difference to provide an indication of any change of symptoms.

#### Data Preprocessing

*Electroencephalogram (EEG).* EEG data was band-pass filtered (2–100 Hz) using a third-order Butterworth filter. Data were manually inspected for artifact (eye blink, muscle activity) and contaminated epochs removed. Two time-windows were then extracted from the data for analysis: A 5-minute resting baseline and the CHEP pain testing (~10 minutes). These time windows are represented in **Figure 1** as the Baseline and CHEP Testing periods. To examine LIFU effects on the CHEP, the data within the CHEP testing time-window was epoched −2000 to 2000 msec around each of the 40 trials and baseline corrected (−1500 to −500 ms). *Electrocardiogram.* ECG data was filtered from 10-30 Hz using a third order Butterworth filter. Data was then manually inspected for movement artifact and removed. R-R peaks were found using the automated detection program *findpeaks* in Matlab and confirmed manually. Data was then divided into the same two epochs as the EEG data (Baseline, CHEP testing).

*Electrodermal Response.* EDR data was zero meaned and filtered using a bandpass (0.1 – 25 Hz) third order Butterworth filter. Data was manually inspected for movement artifact and removed. Data was divided into the same two epochs as the EEG data (Baseline, CHEP testing). Within the CHEP testing window, EDR data was further epoched around the onset of the CHEP stimulus (−5 to 10 seconds), mean subtracted and averaged for the 40 CHEPs trials for each condition for each individual.

### Data Analysis

For all variables of interest, Bartlett’s test was used to assess homoscedasticity. Variables that violated homogeneity of variances were evaluated using non-parametric statistical tests.

#### Pain Ratings

*Group Average Pain Ratings*. Pain ratings were averaged across the 40 trials and participants and compared across conditions (LIFU, Sham). Six participants were missing data for one or more trials for either the Sham or LIFU condition due to either a lack of response or data acquisition error. The maximum number of missing ratings within a single participant was three. To account for this, the appropriate number of ratings were randomly removed from each participant so that each participant had 37 ratings for the group-averaged pain rating analysis. Values are reported as mean ± SEM. A paired t-test was used to compare between conditions at a significance level of p < 0.05.

*Temporal Comparison of Pain Ratings.* We were also interested in the effect of repeated LIFU administration over time and as such performed a temporal analysis across the 40 CHEP trials. Pain ratings for each trial were averaged across participants, creating 40 group-averaged pain ratings for each condition. In the sham condition, one participant had missing data for three trials and two participants were each missing data for one trial. In the LIFU condition, six participants were each missing data for one trial. Missing data was not included in this analysis. The pain rating data was then condensed into 4 bins of 10 pain ratings (ratings 1-10, 11-20, 21-30, and 31-40) for dimensional reduction and statistical comparison. Data was analyzed using a two-way repeated-measures ANOVA with factors CONDITION (LIFU, Sham), TIME (1-10, 11-20, 21-30, 31-40), and TIME*CONDITION interaction. Any significant effects were investigated with post-hoc testing at a significance level of p < 0.05.

#### Homeostatic Metrics

*Heart Rate Variability.* Common heart rate variability (HRV) metrics were calculated according to prior literature (Shaffer and Ginsberg, 2017). Time-domain metrics included the mean difference between normal sinus beats (MNN), the standard deviation of the inter-beat interval for normal sinus beats (SDNN), root mean squared of successive differences (RMSSD), percentage of successive intervals that differ by more than 50ms (pNN50), and the coefficient of variation for successive differences between normal sinus beats (σ/μ) (CV). Time-domain metrics during CHEP testing were normalized to their respective mean of the 5-minute baseline data and compared across Sham and LIFU conditions using a paired t-test at a significance level of p < 0.05. Data is reported as the mean ± SEM normalized units where 1 represents the mean of the 5-minute baseline time window.

For the frequency domain, R-R interval data were cubic spline interpolated and transformed using the fast-fourier transform. Data during CHEP testing was normalized to the 5-minute pre-task baseline data, and the normalized powers were averaged across the LF power (0.04-0.15 Hz) and HF power (0.15-0.4 Hz) bands for each subject for each condition (Shaffer and Ginsberg, 2017). The normalized LF/HF ratio was calculated by dividing the LF by the HF power. Normalized frequency-domain powers were compared across Sham and LIFU conditions using a paired t-test at a significance level of p < 0.05. To investigate changes across specific HRV frequencies with each of the LF and HF bands, we performed non-parametric permutation testing of each frequency across conditions (Sham, LIFU) from 0.04-0.4 Hz. Permutation statistics were generated using 1000 iterations at each frequency interval (0.001 Hz). Results were then cluster thresholded to 10 intervals, requiring a consecutive 0.01 Hz to be significant at p < 0.05.

*Electrodermal Response.* Mean peak-to-peak (absolute of min-max) of the EDR data in the interval 0-10 seconds after the CHEP stimulus were compared using a Wilcoxon signed rank test with significance level of p < 0.05.

*Blood Pressure.* Post-session minus pre-session differences in systolic (SBP) and diastolic (DBP) blood pressures and mean arterial pressure (MAP) were calculated and compared between conditions. MAP was calculated according to the common equation used for BP acquired from a standard blood pressure cuff: 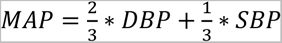 (Razminia et al., 2004). One subject’s arm was unable to fit in the pressure cuff and thus is not included in blood pressure analysis. Values for the 15 included participants are reported as mean ± SEM. Paired t-tests were conducted between conditions at a significance level of p < 0.05.

#### EEG

*CHEP Peak-to-Peak Analysis.* The pre-processed data was first averaged across the 40 trials for each condition and each participant from the time-window −2000ms to +2000ms around the CHEP stimulus. Classically defined CHEP components (De Schoenmacker et al., 2021) were assessed by manually identifying the N1 and P1 peaks in the trial-averaged data for each condition and participant. A distinct inflection of the waveform was necessary for inclusion in statistical analyses. Due to data acquisition errors, several participants were missing data for one or more CHEP stimuli. The maximum number of missing trials within a single participant was two. To account for this, two random trials were removed from each participant, leaving 38 stimuli for each participant for the peak-to-peak analysis. The N1-P1 peak-to-peak amplitude as well as individual N1 and P1 amplitudes were then compared for differences between LIFU and Sham conditions. Wilcoxon signed rank test was used for all CHEP amplitude testing with significance set at p < 0.05.

*Temporal Comparison of CHEP Amplitudes.* Temporal analysis of the CHEP amplitude was conducted to investigate effects across time. Automated detection of minimum and maximum peak was performed for each trial for each participant and condition using the window 200-700ms following the onset of the CHEP stimulus. Two participants were missing the 40^th^ trial for the LIFU condition and one participant was missing the 40^th^ trial for the Sham condition. To account for these three missing trials, the 39^th^ stimulation was duplicated, creating 40 stimulations for all participants. This duplication was done instead of permitting missing data or removing trials so that the data could be equally binned into the 4 averages of 10 stimulations. The CHEP peak-to-peak amplitudes were then averaged across participants, creating 40 group-averaged amplitudes for each condition. The data was then binned into 4 averages of 10 stimulations for statistical comparison. Data was analyzed by performing a two-way repeated-measures ANOVA with factors CONDITION (LIFU, Sham), TIME (trials 1-10, 11-20, 21-30, 31-40), and TIME*CONDITION interaction. Any significant effects within each factor were investigated with post-hoc testing at a significance threshold of p < 0.05.

*Correlation between CHEP Amplitudes and Pain Ratings.* We next aimed to investigate what the relationship was between CHEP amplitudes and pain ratings and if LIFU changed this relationship. To test this, the CHEP amplitudes (N1-P1 peak-to-peak, N1, and P1) and pain ratings were averaged across participants for each of the 40 trials in the same way as above for temporal comparisons, therefore the same approach was taken for missing data. The N1-P1 peak-to-peak CHEP amplitude, the N1 amplitude, and the P1 amplitude were each correlated with pain ratings across trials using Pearson’s correlations for both Sham and LIFU conditions. Rho values were then transformed into z-scores using Fisher’s r-to-z transformation (Meng et al., 1992). Z-scores of the correlation coefficients were then compared between the LIFU and Sham conditions using Fisher’s z-test at a significance level of p < 0.05.

*EEG Time-Frequency*. Morlet wavelet convolution was applied to the pre-processed data using 30 log-spaced frequencies from 2-100 Hz in the time-window −2000ms to +2000ms around the CHEP stimulus at time zero. We then conducted two analyses: a canonical mean frequency band analysis only around the time of the CHEP (200 – 600 ms) and also a broader time-frequency non-parametric permutation analysis to investigate any specific EEG power changes over a wider time period (−100 to 1000 ms).

*Frequency band analysis*. Frequency band analysis was conducted by averaging EEG power for the time period of 200 to 600ms following the onset of the CHEP stimulus and binning the frequencies into their respective canonical bands including delta (2-4 Hz), theta (4-8 Hz), alpha (8-13 Hz), beta (13-30 Hz), low gamma (30-59 Hz), and high gamma (61-100 Hz). The power for each frequency band was baseline corrected to the time window −1500ms to −500ms prior to the onset of each stimulus at time zero. Data was then compared between conditions using paired t-tests at a significance level of p < 0.05.

*Non-parametric permutation analysis*. The time-series data for each log-spaced frequency was averaged across participants in 1ms intervals beginning 100ms prior to the CHEP stimulus to 1000ms after the CHEP stimulus. Data was analyzed using permutation statistics (Maris and Oostenveld, 2007) with 1000 permutations at each time point with a significance level of <0.05. P-values were cluster thresholded to 10ms, requiring 10 consecutive milliseconds within each frequency to have a p-value <0.05 in order to be statistically significant.

#### Quantitative Modeling of Ultrasound Wave Propagation

Simulations were performed using the k-Wave Matlab toolbox (Treeby and Cox, 2010), which uses a pseudo-spectral time domain method to solve discretized wave equations on a spatial grid. Acoustic simulations were performed for each participant. Each participants’ dataset consisted of their structural MRI and CT of the head. CT images were used to construct the acoustic model of the skull, while MR images were used to target LIFU to the dACC coordinates. CT and MR images were first co-registered using custom scripts written in Matlab and the images were then upsampled to a finer resolution for use in the acoustic simulations. The skull was extracted manually using a threshold intensity value and the intracranial space was assumed to be homogenous as ultrasound reflections between soft tissues are small (Mueller et al., 2016). Acoustic parameters were calculated from CT data assuming a linear relationship between skull porosity and the acoustic parameters (Aubry et al., 2003; Marquet et al., 2009).

The computational model of the ultrasound transducer used in simulations was constructed to recreate empirical acoustic pressure maps of focused ultrasound transmitted in an acoustic test tank, similar to previous work (Mueller et al., 2016; Legon et al., 2018a). The modeled ultrasound transducer and waveform was designed to match the transducer and waveform used during pain testing. The shape of the waveform produced by the transducer can be visualized in **Figure 2**. The skull properties used in the model are previously reported by Legon et al., 2018 (Legon et al., 2018a). Estimated intracranial pressure values were extracted from each participants’ dACC brain volume, expressed in kilopascals (kPa) ± (SEM). Intensity parameters were derived from measured values of pressure using the approximation of plane progressive acoustic radiation waves (Preston, 1986).

**Figure 2.**
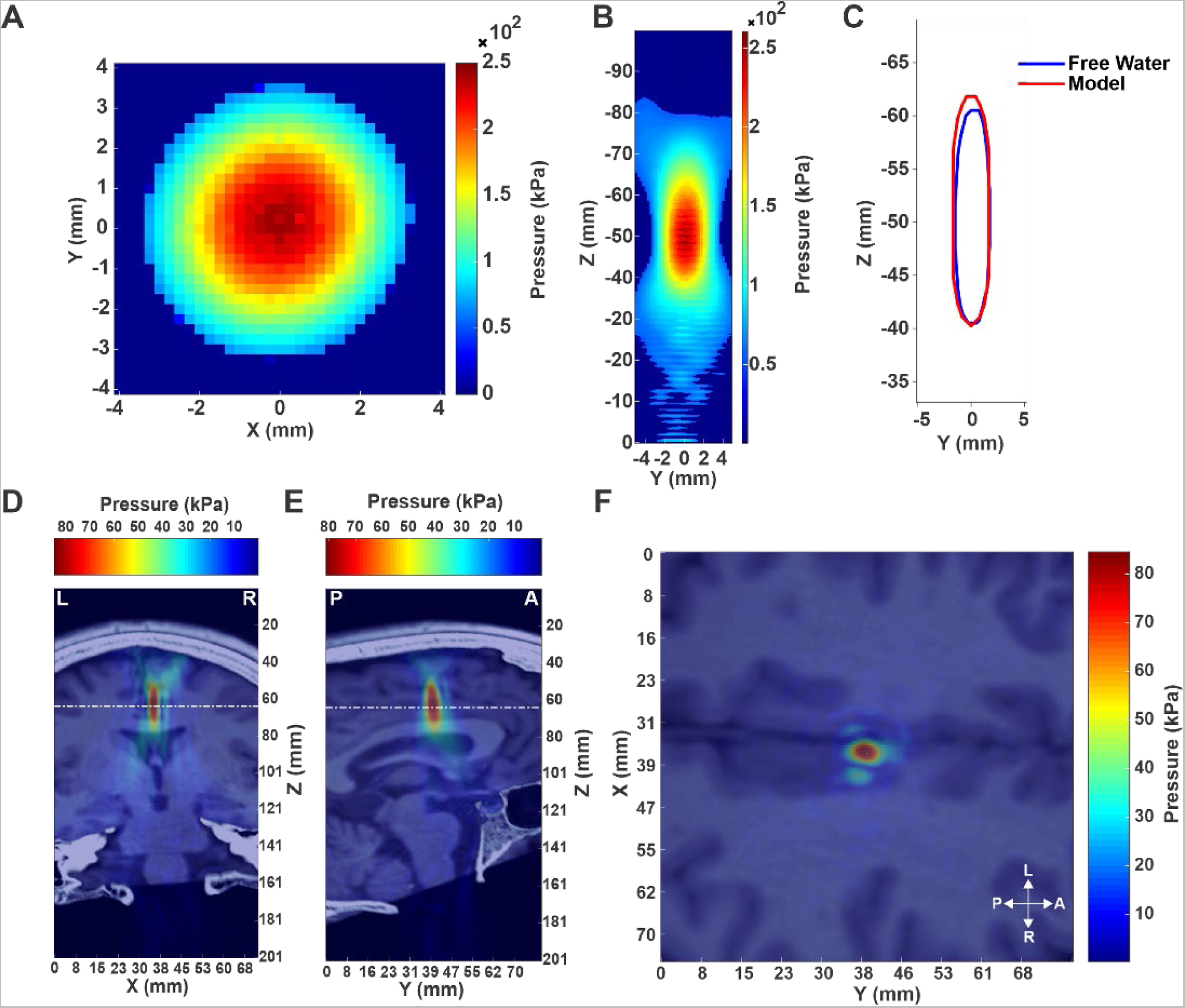
Transducer characteristics and acoustic modelling. **(A)** Pseudo-color free water XY lateral beam pressure profile at Z maximum of the 500kHz single-element transducer measured in kilopascals (kPa). **(B)** Pseudo-color free water YZ axial beam pressure profile. The transducer face is at Z = 0 mm. **(C)** Full-width half maximum (FWHM) overlay of the YZ axial beam profile from the free water tank data (blue) and the modelled transducer (red) used for acoustic modelling. **(D-F)** Acoustic modelling results using individual MRI and CT scans from a single representative subject. **(D)** Coronal (XZ) view depicting left (L) and right (R) directions. **(E)** Sagittal (YZ) view depicting posterior (P) and anterior (A) directions. **(F)** Transverse (XY) cross-section from the white line in the coronal and sagittal views in D and E.

#### LIFU Targeting Data Analysis

The distance from the scalp marker set in the neuronavigation system to the predetermined dACC target was calculated and expressed in millimeters as an indicator of transducer placement (targeting) error. Targeting error was averaged across the 40 trials for each participant in both Sham and LIFU conditions and expressed as mean ± standard deviation. Group-averaged data was compared using a paired t-test to evaluate differences in targeting between conditions at a significance level of p < 0.05.

#### Analysis of Auditory Confounds

Data from the auditory query for each question (all scored from 1-7) were first collapsed into a 1-3 score in order to reduce the dimensions for comparison between conditions. An original response of a 1-3 (“Strongly Disagree”, “Disagree”, “Slightly Disagree”) was given a score of one indicating a response of “Disagree.” A score of four was assigned a value of two indicating a response of “Unsure.” An original score of 5-7 (“Slightly Agree”, “Agree”, and “Strongly Agree”) was assigned a score of three indicating a response of “Agree.” Data was averaged across participants for each condition and compared using a signed rank test at a significance level of p < 0.05.

## RESULTS

### Empirical Acoustic Measurements

Measurements of the transducer’s beam profile in free water revealed a full-width half maximum (FWHM) lateral (XY) resolution of ± 1.5 mm at the axial (Z) beam focus (**Figure 2A**) and a FWHM axial (YZ) resolution of ± 10mm with a focal depth of 52mm from the exit plane of the transducer (**Figure 2B**). The axial FWHM comparison between the empirical measurements in free water and the modelled waveform demonstrated good agreement validating use in the acoustic models (**Figure 2C**).

### Acoustic Modelling

The extracranial peak negative pressure was 370 kPa for all participants. This translated to an intensity outside of the skull of 4.5 W/cm^2^ spatial-peak, pulsed averaged (I_sppa_) or 1.62 W/cm^2^ spatial-peak, temporal averaged (I_spta_). Acoustic modelling estimated a mean ± SD intracranial pressure at the dACC target of 60.13 ± 15.27 kilopascals (kPa) with a range of 37.36 to 86.19 kPa (**Table S1**). Modelling results of the intracranial beam profile in a single representative subject in the coronal (**Figure 2D**), sagittal (**Figure 2E**), and transverse (**Figure 2F**) views show targeting of the dACC. **Table S1** shows individual subject estimated intracranial pressure.

### LIFU Targeting

The mean ± SD depth from the scalp to the dACC target across all participants was 47.34 ± 3.62 mm (range: 41.5 to 54.5 mm). For females the mean ± SD depth was 45.73 ± 2.76 mm (range: 41.5 to 51.6 mm) and for males it was 50.02 ± 3.44 mm (range: 45.6 to 54.5 mm). Given the FWHM of the axial beam profile at ± 10mm and focus at 52 mm, the beam overlapped well with the dACC target across participants. **Table S1** shows individual subject depths of dACC targets from the scalp.

The mean ± SD targeting error across participants was 1.50 ± 0.81 mm in the Sham condition and 1.18 ± 0.40 mm in the LIFU condition (**Figure S1A**). A paired t-test showed no significant difference between conditions (t_(15)_ = −1.39, p = 0.18). A histogram of the distance from the target for each LIFU stimulation across all participants and conditions can be seen in **Figure S1B**.

### Pain Ratings

The mean ± SEM pain ratings were 3.61 ± 0.26 for the Sham condition and 2.52 ± 0.37 for the LIFU condition, with a difference of −1.08 ± 0.21 or −30.2% (range: −2.30 to +0.38) under the LIFU condition relative to Sham. A paired t-test revealed a significant difference in pain ratings between conditions (t_(15)_ = −5.4, p = 7.4×10^−5^) (**Figure 3A**).

**Figure 3.**
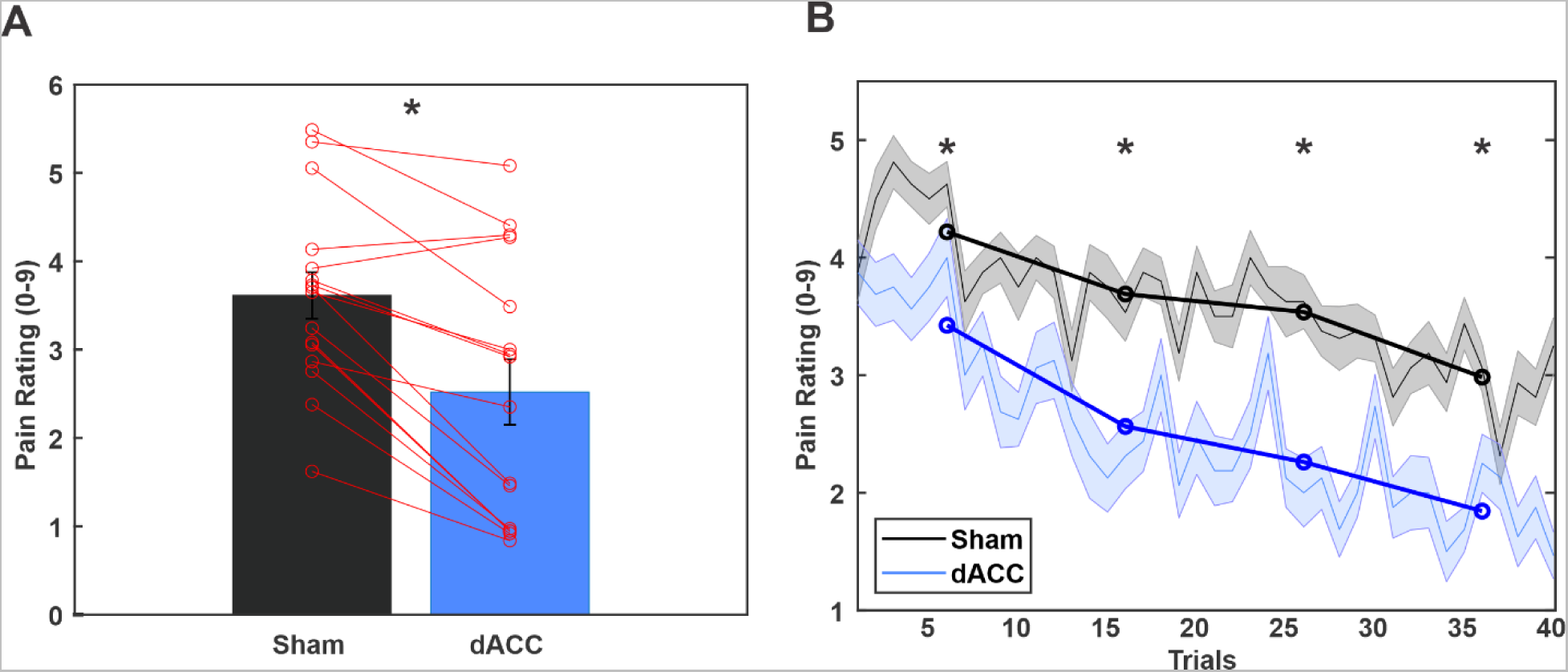
Effect of LIFU to dACC on pain ratings. **(A)** Group (N=16) mean ± SEM pain ratings (0-9 scale). Red lines represent individual subject data. Asterisk indicates significant differences between conditions at p < 0.0001. **(B)** Group (N=16) mean ± SEM pain ratings across 40 trials for Sham (black) and low-intensity focused ultrasound (LIFU) to the dorsal anterior cingulate (dACC) (blue). Thin lines and shaded regions are the mean ± SEM for individual trials. Thick lines represent every 10 trial averages. Asterisks represent significant differences in 10 trial averages across conditions at p < 0.05.

To investigate if pain ratings changed across the 40 trials, we performed a two-way repeated-measures ANOVA with factors TIME (trials 1-10, 11-20, 21-30, 31-40) and CONDITION. Results demonstrated a significant main effect of CONDITION (F_(1,15)_ = 19.88, p = 1.87×10^−5^) and TIME (F_(3,15)_ = 5.79, p = 0.001) but no effect of CONDITION*TIME interaction (F_(3,15)_ = 0.51, p = 0.94). Post hoc testing across conditions for each binned time point revealed LIFU significantly reduced pain ratings at each time bin including trials 1-10 (t_(15)_ = −3.72, p = 0.0021), 11-20 (t_(15)_ = −4.44, p = 4.8×10^−4^), 21-30 (t_(15)_ = −4.57, p = 3.7×10^−4^), and 31-40 (t_(15)_ = −4.20, p = 7.7×10^−4^) (**Figure 3B**).

### Cardiovascular responses

#### Time domain

*SDNN.* The mean ± SEM normalized SDNN for the Sham condition was 0.77 ± 0.044 and for the LIFU condition was 0.95 ± 0.063 with a difference of +0.15 ± 0.043 under the LIFU condition relative to Sham. A paired t-test showed a significant difference between the conditions (t_(15)_ = 3.76, p = 0.002) (**Figure 4A**). This would suggest that during painful stimuli, SDNN is reduced as compared to baseline levels (below 1) and that LIFU looks to return these levels towards baseline resting levels.

**Figure 4.**
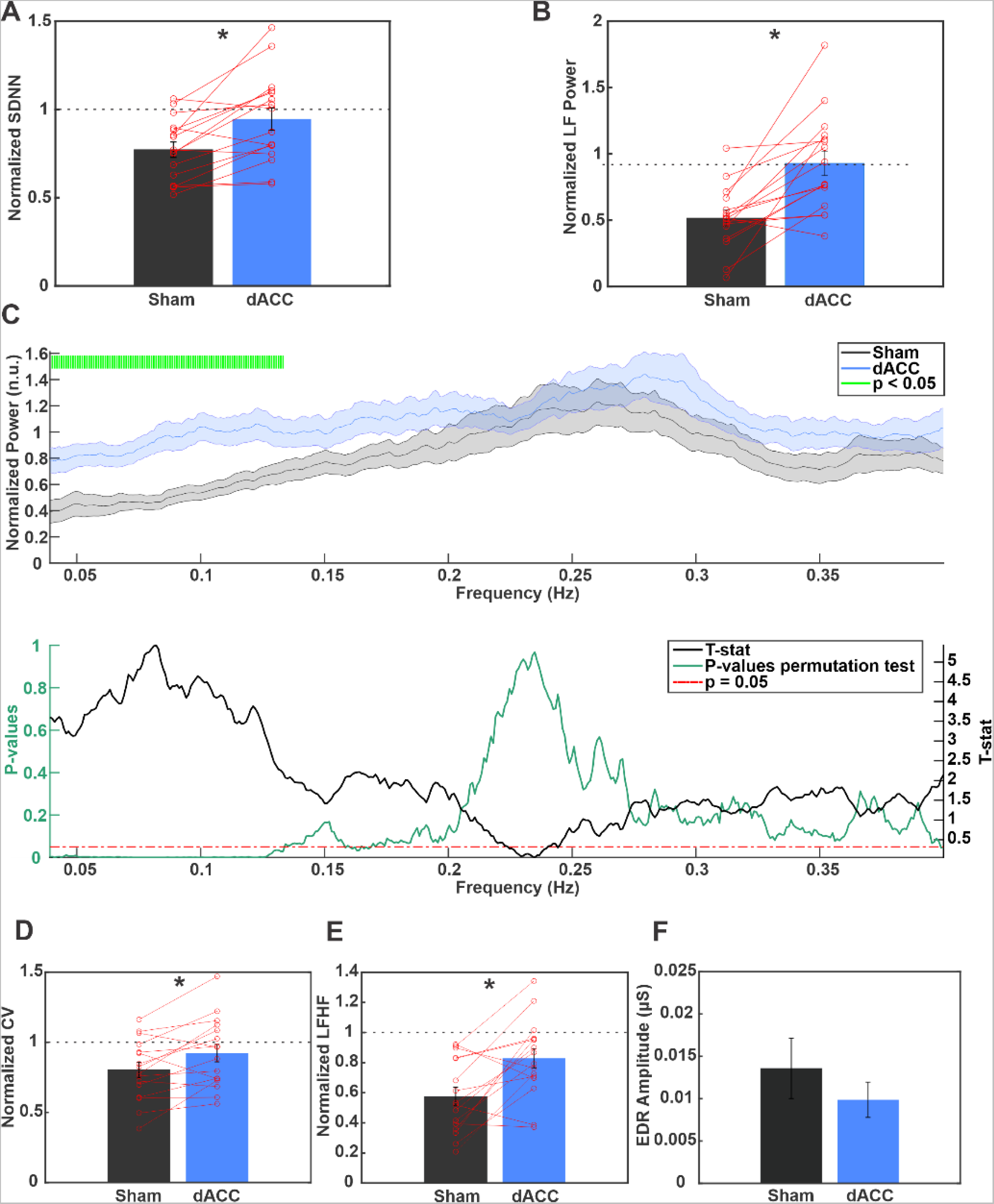
Effect of LIFU to the dACC on homeostatic cardiovascular responses. A, B, D, and E depict measures of heart rate variability derived from electrocardiogram data where the Y-axis is the metric during contact heat-evoked potential (CHEP) testing normalized to the metric during the 5-minute baseline period which is represented by the horizontal dashed line at 1. For these figures, red lines represent individual subject data, and the asterisk indicates significant differences between conditions at p < 0.01. **(A)** Group (N=16) mean ± SEM standard deviation of normal sinus beats (SDNN). **(B)** Group (N=16) mean ± SEM low-frequency (LF) power. **(C)** (Top) Group (N=16) mean ± SEM normalized HRV power. X-axis is frequency in hertz (Hz) and y-axis is power during CHEP testing normalized to power during the 5-minute baseline period. Lines and shaded regions represent the mean ± SEM for sham (black) and low-intensity focused ultrasound (LIFU) to the dorsal anterior cingulate (dACC) (blue). Green highlights represent significant differences between conditions at p<0.05 cluster thresholded to 0.01 Hz. (Bottom) Results of permutation testing. T-statistics (black line) and p-values (green line) from permutation testing across frequencies. The dashed red line represents p = 0.05. P-values that remain below p < 0.05 line up to or beyond the cluster threshold are represented in C with a green highlight. **(D)** Group (N=16) mean ± SEM coefficient of variation (CV). **(E)** Group (N=16) mean ± SEM low-frequency/high-frequency (LFHF) ratio. Y-axis is the LFHF ratio during CHEP testing normalized to the LFHF ratio during the 5-minute baseline period which is represented by the horizontal dashed line at 1. **(F)** Group (N=16) mean ± SEM electrodermal response (EDR). Y-axis is EDR amplitude in microsiemens (μS).

*Coefficient of variation (CV)*. The mean ± SEM normalized Coefficient of Variation (CV) was 0.80 ± 0.054 for the Sham condition and 0.92 ± 0.06 for the LIFU condition with a difference of +0.12 ± 0.04 under the LIFU condition relative to Sham. A paired t-test demonstrated a statistically significant difference between conditions (t_(15)_ = 3.01, p = 0.009) (**Figure 4D**).

*MNN*. The mean ± SEM normalized MNN was 1.03 ± 0.02 under the Sham condition and 1.03 ± 0.01 under the LIFU condition with a difference of 0.0013 ± 0.014. A paired t-test showed no significant difference between conditions (t_(15)_ = −0.09, p = 0.93).

*RMSS*D. The mean ± SEM normalized RMSSD was 1.03 ± 0.02 under the Sham condition and 1.03 ± 0.01 under the LIFU condition with a difference of 0.0002 ± 0.014. A paired t-test showed no significant difference between conditions (t_(15)_ = −0.01, p = 0.99).

*pNN50.* The mean ± SEM normalized pNN50 was 1.38 ± 0.16 under the Sham condition and 1.41 ± 0.12 under the LIFU condition for a difference of 0.031 ± 0.22. A paired t-test showed no significant difference between conditions (t_(15)_ = 0.14, p = 0.89) (**Figure S5**).

#### Frequency domain

*Low-frequency (LF) power.* The mean ± SEM normalized LF power for the Sham condition was 0.52 ± 0.06 and 0.93 ± 0.09 for the LIFU condition with a difference of +0.41 ± 0.09 under the LIFU condition relative to Sham. A paired t-test demonstrated a significant difference between conditions (t_(15)_ = 4.54, p < 0.001) (**Figure 4B**).

*High-frequency (HF) power*. The mean ± SEM normalized HF power for the Sham condition was 0.95 ± 0.11 and for the LIFU condition was 1.13 ± 0.01 with a difference of +0.18 ± 0.10 under the LIFU condition relative to Sham. A paired t-test showed no significant difference between conditions (t_(15)_ = 1.79, p = 0.09) (**Figure S5**).

*Non-parametric permutation testing of frequency.* We followed the analysis of mean normalized LF and HF powers with permutation testing (1000 permutations; p <0.05) to investigate the effects of LIFU across HRV powers from 0.04-0.15 Hz. Permutation statistics revealed LIFU to the dACC increased normalized LF power in the frequency range 0.04-0.13Hz. There were no significant differences in normalized HF power in the permutation testing (**Figure 4C**).

*LF/HF Ratio.* The mean ± SEM normalized LF/HF ratio in the Sham condition was 0.57 ± 0.06 and 0.83 ± 0.06 in the LIFU condition with a difference of +0.26 ± 0.07 under the LIFU condition relative to Sham. A paired t-test demonstrated a significant difference between conditions (t_(15)_ = 3.61, p = 0.003) (**Figure 4E**).

*Electrodermal Response.* EDR mean ± SEM amplitude was 0.014 ± 0.0036 μS for the Sham condition and 0.009 ± 0.0021 μS for the LIFU condition. The Wilcoxon sign rank revealed a non-significant difference of −0.0037 ± 0.0031 μS between conditions (z = −1.31, p = 0.19) (**Figure 4F**). *Blood Pressure.* Under the Sham condition, the mean ± SEM pre-session SBP was 119.27 ± 2.84 mmHg and the DBP was 70.33 ± 1.86 mmHg. The mean ± SEM post-session SBP was 119.00 ± 2.55 mmHg and the DBP was 69.6 ± 2.34 mmHg. Under the LIFU condition, the mean ± SEM pre-session SBP was 121.2 ± 3.21 mmHg and the DBP was 70.87 ± 2.15 mmHg. The mean ± SEM post-session SBP was 117.67 ± 3.75 mmHg and the DBP was 71.27 ± 2.68 mmHg. Mean ± SEM post-session minus pre-session SBP differences were −0.27 ± 2.57 mmHg for Sham and −3.53 ± 1.89 mmHg for LIFU. A paired t-test revealed no significant difference between conditions (t_(14)_ = −1.29, p = 0.22). Mean ± SEM post-session minus pre-session DBP differences were −0.73 ± 1.42 mmHg for Sham and +0.40 ± 1.27 mmHg for LIFU. A paired t-test showed no significant difference between conditions (t_(14)_ = 1.03, p = 0.32).

*Mean Arterial Pressure (MAP).* Under the Sham condition, the mean ± SEM pre-session MAP was 86.64 ± 2.07 while post-session MAP was 87.64 ± 2.31. Under the LIFU condition, the mean ± SEM pre-session MAP was 86.07 ± 2.28 while the post-session MAP was 86.73 ± 2.84. The mean ± SEM post-session minus pre-session MAP differences were −0.58 ± 1.57 mmHg for Sham and −0.91 ± 1.25 mmHg for LIFU. A paired t-test revealed no significant difference between conditions (t_(14)_ = −0.23, p = 0.82).

### CHEP Amplitude

We conducted peak-to-peak analysis to test whether LIFU affected the N1-P1 peak-to-peak amplitude of the CHEP as well as the N1 and P1 amplitudes separately. The mean ± SEM N1-P1 peak-to-peak amplitude was 26.73 ± 0.73 μV for the Sham condition and 20.06 ± 0.48 μV for the LIFU condition for a difference of −6.67 ± 2.81 μV or −25.1% under the LIFU condition relative to Sham. A Wilcoxon signed rank test revealed a significant difference in the mean N1-P1 amplitude between conditions (z = −2.17, p = 0.030) (**Figure 5A & C**).

**Figure 5.**
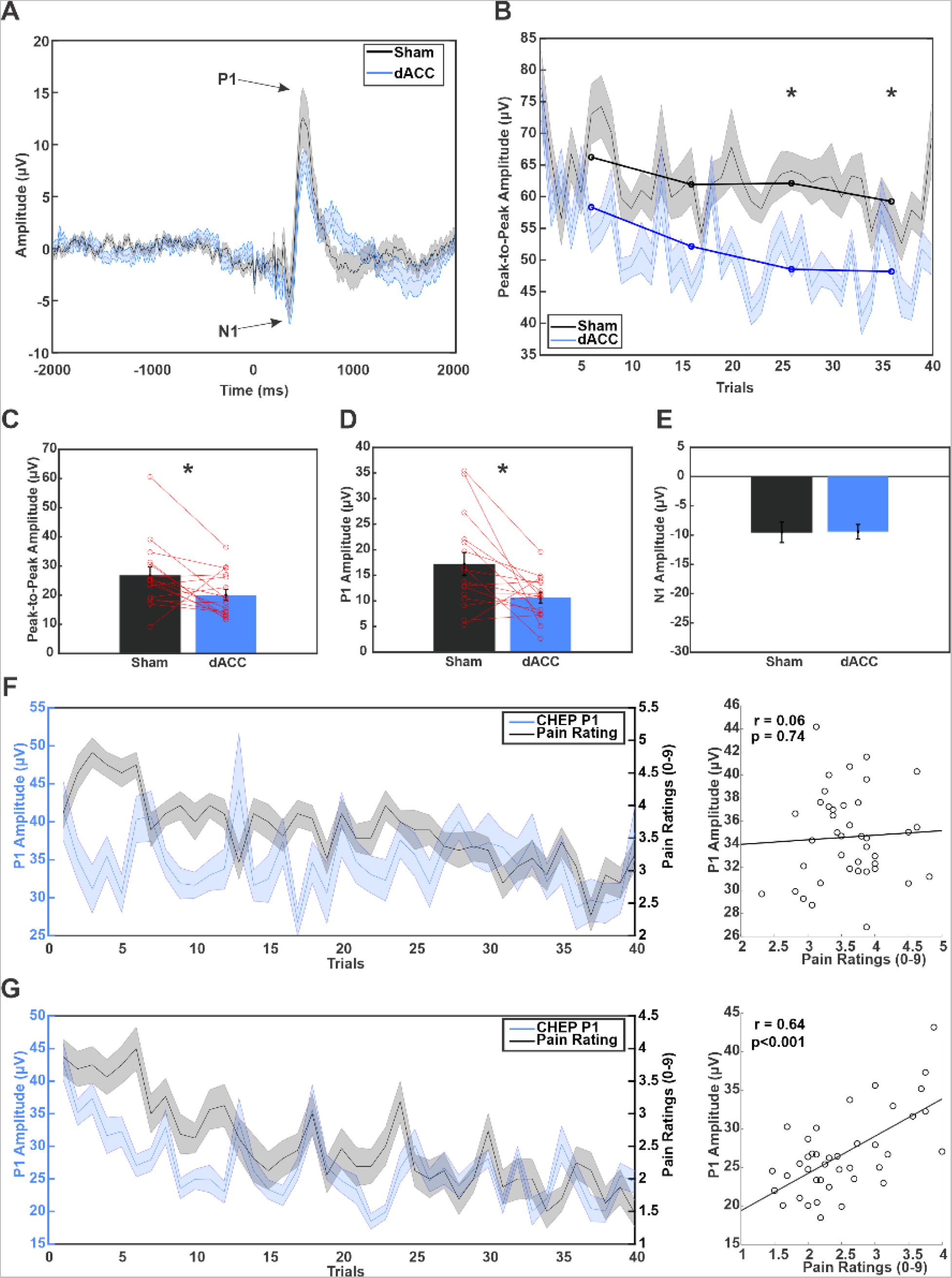
Effect of LIFU to the dACC on CHEP amplitudes and behavior. **(A)** Group (N=16) mean ± SEM contact heat-evoked potential (CHEP) trace for sham (black) and low-intensity focused ultrasound (LIFU) to the dorsal anterior cingulate (dACC) (blue). Amplitude is in microvolts (μV). Time 0 ms represents delivery of CHEP stimulus to the dorsum of the right hand. Arrows point to the peaks of the N1 and P1 amplitudes. **(B)** Group (N=16) mean ± SEM N1-P1 peak-to-peak CHEP amplitude across the 40 trials for sham (black) and LIFU to dACC (blue) conditions. Y-axis is the peak-to-peak amplitude (μV). Thin lines and shaded regions are the mean ± SEM for each of the 40 trials. Thick lines and dots represent 10 trial averages. Asterisks represent significant differences in 10 trial averages across conditions at p < 0.05. **(C)** Group (N=16) mean ± SEM N1-P1 peak-to-peak CHEP amplitude. Red lines represent individual subject data. Asterisk denotes a significant difference between groups at p < 0.05. **(D)** Group (N=16) mean ± SEM P1 CHEP amplitude for each condition. Red lines represent individual subject data. Asterisk denotes a significant difference between groups at p < 0.05. **(E)** Group (N=16) mean ± SEM N1 CHEP amplitude for Sham and LIFU to dACC. **(F)** (Left) Group (N=16) mean ± SEM P1 amplitudes in blue and pain ratings in black across the 40 trials for the Sham condition. (Right) Group (N=16) scatter plot depicting the correlation between the P1 amplitude and pain ratings across trials for the Sham condition. r = correlation coefficient, p = p-value, black line = least-squares line. **(G)** (left) Group (N=16) mean ± SEM P1 amplitudes in blue and pain ratings in black across the 40 trials for the LIFU to dACC condition. (right) Group (N=16) scatter plot depicting the correlation between the P1 amplitude and pain ratings across trials for the LIFU to dACC condition. r = correlation coefficient, p = p-value, black line = least-squares line.

Investigation of the N1 and P1 components of the CHEP separately demonstrated the mean ± SEM amplitude of the P1 component was 17.20 ± 2.28 μV for the Sham condition and 10.65 ± 1.05 μV for the LIFU condition with a difference of −6.54 ± 2.39 μV or −38.1% under the LIFU condition relative to Sham. The Wilcoxon signed rank test showed a significant difference in the P1 component between conditions (z = −2.48, p = 0.013) (**Figure 5D**). The mean ± SEM amplitude of the N1 component was 9.53 ± 1.74 μV for the Sham condition and 9.41 ± 1.23 μV for the LIFU condition. The Wilcoxon signed rank test showed no significant difference in the N1 component between conditions (z = −0.05, p = 0.96) (**Figure 5E**).

We also investigated the effect of LIFU on the CHEP over the 40 trials. To do so, we performed a two-way repeated-measures ANOVA with factors TIME (trials 1-10, 11-20, 21-30, 31-40) and CONDITION. Results demonstrated a significant main effect of CONDITION (F_(1,15)_ = 16.47, p = 8.84×10^−5^) and TIME (F_(3,15)_ = 3.28, p = 0.023) but no CONDITION*TIME interaction (F_(3,15)_ = 0.31, p = 0.82). Post hoc testing across conditions for each binned time point revealed LIFU to reduce the amplitude of the N1-P1 CHEP in trials 21-30 (t_(15)_ = −3.28, p = 0.005) and 31-40 (t_(15)_ = −2.47, p = 0.026) but not trials 1-10 (t_(15)_ = −1.96, p = 0.068) or 11-20 (t_(15)_ = −2.05, p = 0.059) (**Figure 5B**).

### CHEP Amplitude and Pain Rating Correlations

To examine the relationship between CHEP amplitudes and behavior, we performed correlations between pain ratings and the N1-P1 peak-to-peak amplitude, N1 amplitude, and P1 amplitude across the 40 trials for each condition. We used Fisher’s Z test to analyze the change in correlations between conditions (Meng et al., 1992). The N1-P1 peak-to-peak amplitude did not significantly correlate with pain ratings for the Sham condition (r = 0.31, p = 0.054) but showed a significant positive correlation for the LIFU condition (r = 0.64, p=8.17×10^−6^). Comparison of the fisher z-transformed correlation coefficients revealed a non-significant increase in the correlation under the LIFU condition (z = 1.91, p = 0.057) (**Figure S2A & B**). The N1 amplitude had a significant positive correlation with pain ratings for the Sham condition (r = 0.42, p=0.007) and for the LIFU condition (r = 0.43, p=0.005). Comparison of the fisher z-transformed correlation coefficients revealed no significant difference between conditions (z = −0.06, p = 0.95) (**Figure S2C & D**). The P1 component did not correlate with pain ratings for the Sham condition (r = 0.06, p = 0.74) but had a significant positive correlation for the LIFU condition (r = 0.64, p = 9.64×10^−6^) (**Figure 5F & G**). Comparison of the fisher z-transformed correlation coefficients revealed a significantly increased correlation for the LIFU condition (z = 3.01, p = 0.003).

### EEG Time-Frequency Analysis

We analyzed EEG power by first grouping the data into canonical frequency bands and looking at group averaged normalized power from +200 to +600ms after the heat stimulus to investigate power changes only at the time of the CHEP. *Delta (2-4 Hz).* The mean ± SEM normalized power in the delta frequencies was 1.44 ± 0.056 in the Sham condition and 1.22 ± 0.040 in the LIFU condition for a decrease of 0.21 ± 0.076 under the LIFU condition relative to Sham. A paired t-test showed a significant difference between conditions (t_(15)_ = −2.79, p = 0.014).

*Theta (4-8 Hz).* The mean ± SEM normalized power in the theta frequencies was 1.35 ± 0.045 in the Sham condition and 1.23 ± 0.033 in the LIFU condition for a decrease of 0.12 ± 0.059 under the LIFU condition relative to Sham. A paired t-test showed no significant difference between conditions (t_(15)_ = −1.99, p = 0.065).

*Alpha (8-12 Hz).* The mean ± SEM normalized power in the alpha frequencies was 1.15 ± 0.029 in the Sham condition and 1.08 ± 0.032 in the LIFU condition for a decrease of 0.07 ± 0.042 under the LIFU condition relative to Sham. A paired t-test showed no significant difference between conditions (t_(15)_ = −1.75, p = 0.10).

*Beta (13-30 Hz).* The mean ± SEM normalized power in the beta frequencies was 1.02 ± 0.022 in the Sham condition and 0.99 ± 0.020 in the LIFU condition for a decrease of 0.035 ± 0.029 under the LIFU condition relative to Sham. A paired t-test showed no significant difference between conditions (t_(15)_ = −1.21, p = 0.24).

*Low gamma (30-60 Hz).* The mean ± SEM normalized power in the low gamma frequencies was 0.99 ± 0.020 in the Sham condition and 0.98 ± 0.027 in the LIFU condition for a decrease of 0.01 ± 0.026 under the LIFU condition relative to Sham. A paired t-test showed no significant difference between conditions (t_(15)_ = −0.35, p = 0.73).

*High gamma (60-100 Hz).* The mean ± SEM normalized power in the high gamma frequencies was 0.95 ± 0.01 in the Sham condition and 0.97 ± 0.023 in the LIFU condition for an increase of 0.014 ± 0.023 under the LIFU condition relative to Sham. A paired t-test showed no significant difference between conditions (t_(15)_ = 0.61, p = 0.55). Bar plots of the data for all frequency bands can be visualized in **Figure S4**.

We followed the mean frequency band analysis at the time of the CHEP with time-frequency analysis for the time window 100ms prior to 1000ms after the CHEP stimulus to investigate the effects of LIFU over a broader period around the CHEP and for each individual frequency (2 – 100 Hz). Group pseudo-colored time-frequency spectra for the Sham and LIFU conditions can be seen in **Figure 6A** with the LIFU minus Sham difference map in **Figure 6B**. Permutation statistics (1000 permutations; p < 0.05) demonstrated significant decreases in delta power (2-4 Hz) beginning 200ms after the onset of the heat stimulus and ending 800ms post stimulus. Decreases in theta power (4-8 Hz) were observed from 200ms to 400ms after the CHEP stimulus and decreases in beta power (13-30 Hz) around 200 and 300ms after the stimulus. Increases in high gamma power (60-100 Hz) were seen at 100ms and from around 400ms to 1000ms following the stimulus (**Figure 6C**). The raw p-value and thresholded maps can be seen in **Figure S3**.

**Figure 6.**
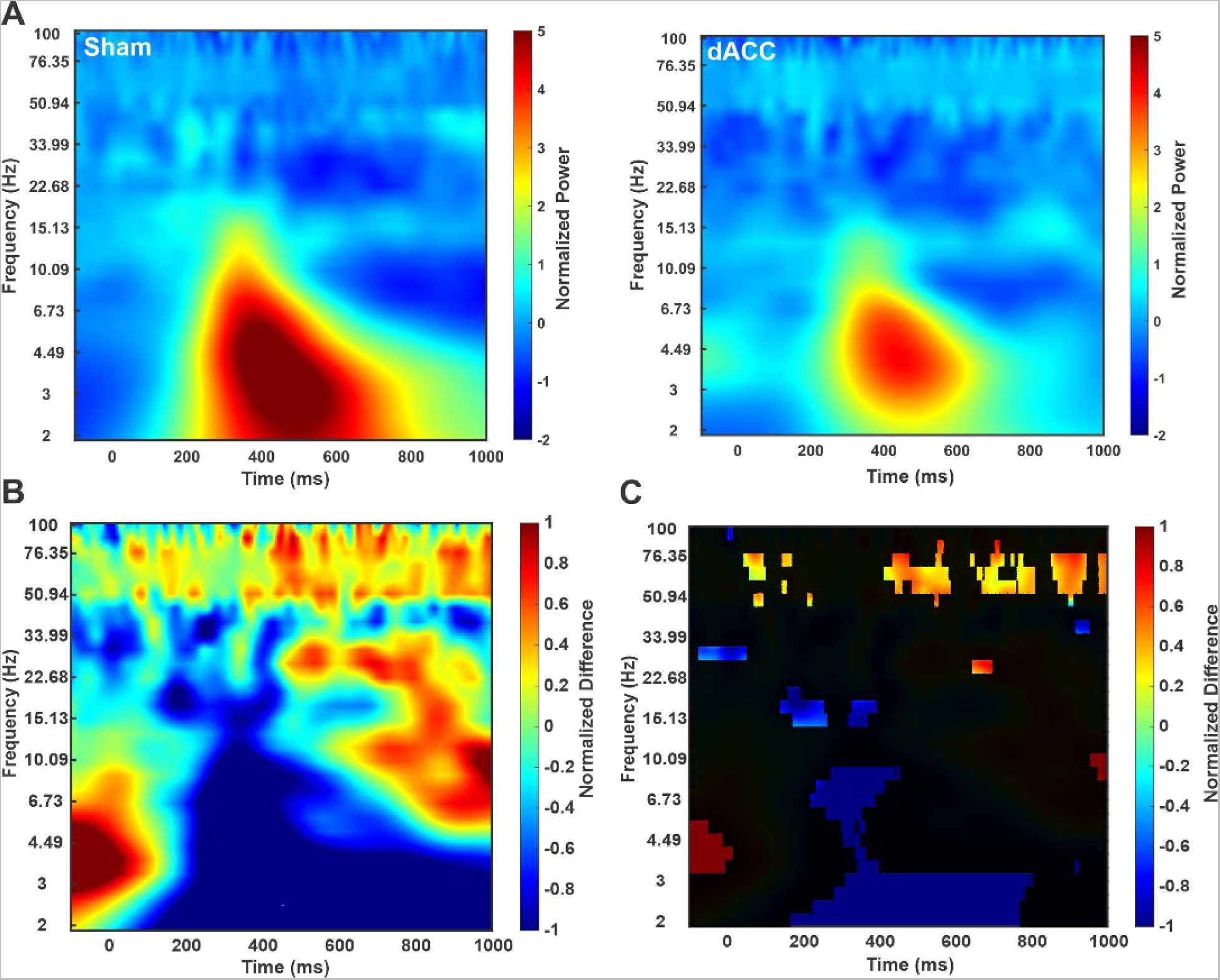
CHEP EEG time-frequency analysis. **(A)** Group (N=16) pseudo-colored time-frequency map from the time window −100ms to 1000ms around the CHEP stimulus (0 ms) during the Sham condition (left) and the low-intensity focused ultrasound (LIFU) to the dorsal anterior cingulate (dACC) condition (right). X-axis is time in milliseconds (ms) and y-axis is frequency in Hertz (Hz). Warmer colors represent increases and cooler colors represent decreases in power normalized to stimulus baseline window. **(B)** Group (N=16) difference map of Figure 5A above (LIFU-Sham). **(C)** Group (N=16) results from permutation testing overlaid on the difference map from B. Areas not statistically significant are in black. All other areas demonstrated a statistically significant difference between Sham and LIFU using permutation statistics (N = 1000; cluster threshold = 10ms; p < 0.05).

### Auditory Masking

*Hearing.* For the question “I could hear the LIFU stimulation,” mean ± SD response for the Sham condition was 1.63 ± 0.89 and 1.79 ± 0.95 for the LIFU condition indicating that participants were between “Disagree” and “Unsure” in their responses. A Wilcoxon sign rank test demonstrated no significant difference between conditions (z = 0.44, p = 0.66).

*Believing.* For the question “I believe I experienced LIFU stimulation,” the mean ± SD response for the Sham condition was 1.31 ± 0.60 and 1.63 ± 0.96 for the LIFU condition. A Wilcoxon sign rank test demonstrated no significant difference between conditions (z = 1.39, p = 0.16).

*Feeling.* For the question “I could feel the LIFU stimulation,” the mean ± SD response for the Sham condition was 1.11 ± 0.46 and 1.05 ± 0.22 for the LIFU condition. A Wilcoxon sign rank test demonstrated no significant difference between conditions (z = −1.00, p = 0.32). Individual subject results for all questions can be seen in **Figure S6**.

### Adverse Events

For the LIFU condition, one participant reported a mild headache, one reported mild unusual scalp sensations, two reported mild neck pain, one reported mild tooth pain, and one reported mild nausea (**Figure S7A**). For the Sham condition, one participant reported mild unusual feelings, one reported mild tingling, one reported mild itching, one reported mild sleepiness, two reported mild inattentiveness, one reported mild twitching, one reported a mild anxious feeling, and three reported mild balance issues (**Figure S7B**). No moderate or serious adverse events were reported for either condition. A breakdown of symptomology is presented in **Figure S7B** for both LIFU and Sham interventions.

## DISCUSSION

We evaluated the effect of single element 500 kHz LIFU to the dACC on homeostatic signals including pain perception, HRV, blood pressure and EDR, and contact heat-evoked potentials during application of brief, noxious contact heat to the dorsum of the right hand. Results indicate LIFU to the dACC reduced pain perception by 1.08 points on a 9-point scale and increased HRV metrics including the SDNN, CV, LF power and LF/HF ratio as compared to Sham stimulation. LIFU attenuated the N1-P1 peak-to-peak CHEP amplitude driven by decreases in P1. Time-frequency analysis revealed significant reductions in delta, theta and beta power and increases in gamma power under the LIFU condition compared to Sham stimulation.

### LIFU to reduce pain perception

Decreases in pain levels by roughly one point or ~ 30% is within the range of pain relief considered clinically meaningful (Rowbotham, 2001; Cepeda et al., 2003; Smith et al., 2020). These results are encouraging given other options for dACC targeting. Deep brain stimulation, an invasive neurosurgical procedure, has been targeted to the dACC for chronic pain with mixed results (Russo and Sheth, 2015). In a recent case series of 24 patients, average pain intensity reduced by 45% (range: −100% to +25%) at 6-month follow-up, however 6 patients experienced the onset of seizures (Boccard et al., 2017). One recent study attempting non-invasive pain modulation using deep TMS to the ACC showed no reductions in pain ratings (Galhardoni et al., 2019). It is likely no effects were observed as electric fields from this TMS coil do not penetrate deep enough nor is selective enough (Deng et al., 2013b; Drakaki et al., 2022) to reach specific dACC targets as used here. We were able to use precise targets (MNI 0,18,30) based upon large meta-analytic datasets to precisely target a small region within the dACC that looks to be common to pain perception and likely accounts for the effects seen here as compared to those with TMS.

### LIFU for Non-invasive Homeostatic Modulation

Brief, painful stimuli elevate heart rate and alter HRV consistent with increased sympathetic or decreased parasympathetic activity like decreased SDNN and HF power (Loggia et al., 2011; Meeuse et al., 2013; Forte et al., 2022). LIFU to the dACC attenuated homeostatic cardiovascular responses to painful stimuli indicated by increases in the SDNN, CV, LF power and LF/HF ratio towards pre-task baseline levels.

The dACC is innervated by spinothalamic input from the ventrocaudal mediodorsal (MDvc) nucleus of the thalamus, which receives bilateral input from parabrachial nucleus andperiaqueductal grey and projects exclusively to the fundus of the cingulate sulcus, providing an integrated homeostatic pathway to the dACC (Ray and Price, 1993; Craig, 2015). The dACC is also considered a homeostatic effector (or output) region due to its direct role in autonomic and behavioral control (Devinsky et al., 1995; Craig, 2002, 2009; Beissner et al., 2013). Direct electrical stimulation (excitation) of the dACC elicits sympathetic autonomic responses including tachycardia and heat sensations (Parvizi et al., 2013; Oane et al., 2020; Xue et al., 2023). Our findings of increased HRV and decreased heat pain sensation support LIFU confers an inhibitory response as it directly opposes the effects of electrical stimulation. It is unclear whether LIFU to dACC altered homeostatic input or output or whether the effects came from direct inhibition of sympathetic activity or some mechanism of increased parasympathetic control.

HRV is a way of measuring homeostatic regulation as it indexes the ability of the autonomic nervous system to dynamically respond to task demands (Rajendra Acharya et al., 2006; ChuDuc et al., 2013). The SDNN is a validated time-domain metric of overall HRV and in short-term recordings, like the present study, SDNN is primarily influenced by parasympathetic activity with a higher SDNN being consistent with relatively reduced sympathetic or increased parasympathetic activity (Shaffer et al., 2014; Shaffer and Ginsberg, 2017). The reduction of SDNN with delivery of pain and the increase towards baseline under the LIFU condition is consistent with direct inhibition of sympathetic activity or indirect facilitation of parasympathetic control and a strong indication we can shift homeostatic responses to painful stimuli. The LF power band (0.04-0.15 Hz) is jointly influenced by the sympathetic and parasympathetic system and correlates with SDNN (Shaffer et al., 2014; Shaffer and Ginsberg, 2017). In resting conditions, such as our 5-minute baseline period, LF power reflects baroreflex/parasympathetic activity, not cardiac sympathetic innervation (Goldstein et al., 2011; Reyes Del Paso et al., 2013). The increased LF power under the LIFU condition thus fits our findings of increased SDNN, providing confirmation using frequency domain HRV metrics. Furthermore, our dACC target for LIFU delivery closely matches coordinates in prior literature on dACC activity and LF power (Critchley et al., 2003; Rebollo et al., 2018). Prior interpretations of the LF/HF ratio primarily indexing sympathovagal balance is controversial (Billman, 2013), and it may be appropriate to interpret the LF/HF ratio based on the influence of measurement conditions on LF power (Shaffer et al., 2014; Shaffer and Ginsberg, 2017).

Taken together, our results point towards LIFU to the dACC either directly or indirectly attenuating sympathetic responses to heat pain as all autonomic metrics like LF power and SDNN decreased during CHEP testing under the Sham condition and increased back towards pre-task baseline levels under the LIFU condition. This is likely through inhibitory mechanisms based upon previous LIFU data (Legon et al., 2014, 2018a, 2023) and the effects on the amplitude of the CHEP (see below). Reductions in sympathetic (or facilitation of parasympathetic) activity may have applications for chronic pain and anxiety disorders as these populations exhibit parasympathetic dysregulation, sympathetic hyper-reactivity, and reduced HRV (Chalmers et al., 2014; Tracy et al., 2016; Zhuo, 2016; Teed et al., 2022).

### The dACC and pain saliency

The dACC is a critical node in the salience network (SN), responsible for orienting attention to novel stimuli and performing dynamic switching between other large-scale brain networks to respond to homeostatic demands (Menon and Uddin, 2010; Menon, 2015; Seeley, 2019). While we did not explicitly test LIFU’s effect on the saliency of pain, our time-frequency analysis demonstrated changes in gamma band oscillations which are a neural correlate of the attentional effects of pain and function to synchronize brain activity in response to pain (Tiemann et al., 2010; Ploner et al., 2017). Gamma band oscillations are related to pain perception in healthy controls, and sourcing of the gamma oscillations include the dACC (Bassez et al., 2020). A recent hypothesis further suggests oscillations in heart rate may relate to changing oscillations in the brain, including the gamma band, and this interaction can change functional connectivity in brain networks associated with salience detection and emotional regulation (Mather and Thayer, 2018). In our study, LIFU to the dACC increased gamma power while reducing pain perception and increasing HRV, suggesting it may be altering the saliency of pain by changing the synchronization of brain responses to pain.

### The dACC Generates Positive Nociceptive Potentials

Peak-to-peak analysis showed LIFU to the dACC reduced the N1-P1 peak-to-peak amplitude of the CHEP driven exclusively by reductions in P1. Analysis of the correlations between CHEP amplitudes and pain ratings showed a significant change in the relationship between the P1 amplitude and pain ratings, but not N1P1 or N1, further suggesting we selectively modulated the P1 amplitude.

Intracranial EEG responses to laser stimuli show the dACC predominately encodes the positive amplitude of the LEP (Bastuji et al., 2016). Dipole sourcing corroborates direct involvement of the dACC in generating the LEP (Jin et al., 2018) while regions of peak cerebral blood flow during LEP stimuli directly matches where we delivered LIFU (Garcia-Larrea et al., 2003).

While it is clear the dACC at least partially contributes to pain evoked potentials, it is unclear precisely what N1 and P1 components of the LEP/CHEP index. Evidence points to attentional modulation of the positive (P1) component of the CHEP with a recent study in autism spectrum disorder showing the P1 strongly correlates with attention to pain, supported by prior LEP work (Ohara et al., 2004; Boyle et al., 2008; Chien et al., 2017). Research using fMRI shows reduced dACC activity during distraction from pain (Bantick et al., 2002). Distraction induced reductions in positive heat-evoked amplitudes appears to be concomitant with reduced unpleasantness ratings, or affective pain (Boyle et al., 2008). That this positive amplitude has been associated with attentional and affective components of pain matches proposed roles of the dACC in pain processing (Rainville et al., 1997; Price, 2000; Shackman et al., 2011; Xiao and Zhang, 2018). The dACC is thus a likely contributor to positive heat-evoked amplitudes while playing a central role in affective and attentional aspects of pain.

### Auditory Masking

LIFU is known to produce auditory effects that can confound observed neuromodulatory effects (Guo et al., 2018; Sato et al., 2018; Johnstone et al., 2021). In a recent study, a multitone random auditory stimulus could effectively mask the sound of LIFU at the same 1 kHz pulse repetition frequency used in the present study (Liang et al., 2023). Using a similar mask, our results demonstrate that subjects did not appreciably hear the LIFU and were unsure if they received LIFU also mitigating potential placebo effects.

## CONCLUSIONS & FUTURE WORK

Single-element 500 kHz LIFU is an effective method to target the dACC to reduce acute pain perception and homeostatic cardiovascular responses. The observed reductions in pain perception approach clinically meaningful magnitudes and the reductions in homeostatic cardiovascular responses suggest LIFU may be an effective tool to attenuate homeostatic signals in the context of pain. Evidence from both the literature and the present study thus points to the dACC as a region that integrates ascending homeostatic information to respond to noxious stimuli. Furthermore, the results point towards the P1 as at least partially generated in the dACC; that LIFU to the dACC may attenuate the affective and attentional component pain in addition to pain intensity, and that LIFU may affect the saliency of acute heat pain. Our findings of reduced pain perception, homeostatic responses, and CHEP amplitudes, for the first time in humans, support prior findings on homeostasis and pain processing through selective, non-invasive, causal manipulation of the dACC.

Future work should aim to evaluate the longevity and durability of LIFU’s effects on these measures as well as its efficacy as a therapeutic option in chronic pain populations. Additionally, it should test the ability of LIFU to the dACC to differentially impact pain intensity, affective pain, and pain saliency.

## Acknowledgements

This work was funded in part by grants awarded to WL from the Seale Innovation Fund and the Focused Ultrasound Foundation. The authors would like to thank Jessica Florig for help with data collection and analysis.

## SUPPLEMENTAL FIGURES

**Figure S1.**
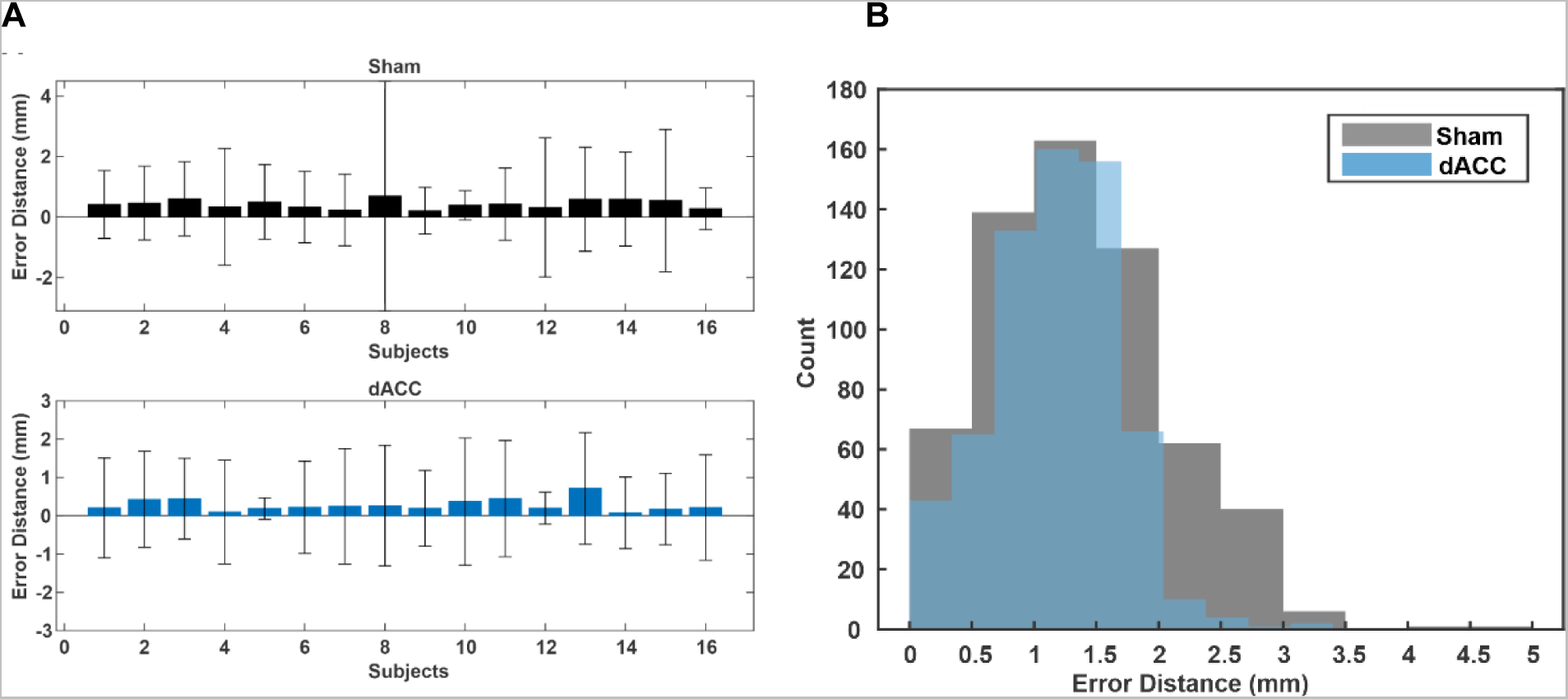
Targeting error. **(A)** (Top) Mean ± standard deviation (SD) targeting error distance of the transducer on the scalp in millimeters (mm) for each subject in the sham condition. Subject 8 in the sham condition shows high variability due to loss of data. (Bottom) Mean ± SD targeting error distance in millimeters (mm) for each subject in the low-intensity focused ultrasound (LIFU) to the dorsal anterior cingulate (dACC) condition. **(B)** Group (N=16) histogram of the absolute targeting error (mm) for both Sham (black) and LIFU to dACC (blue) conditions. X-axis is targeting error in 0.5mm bins.

**Figure S2.**
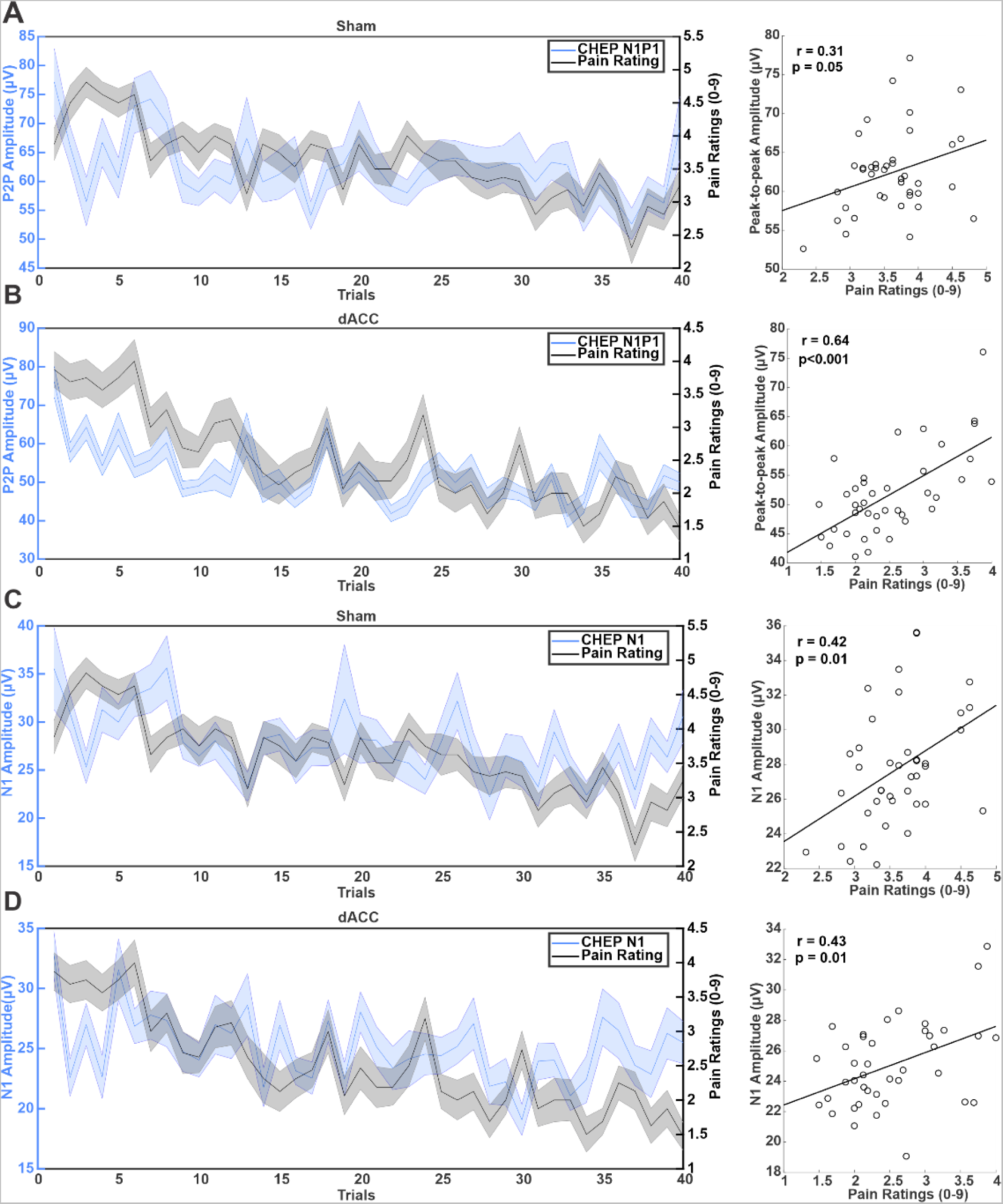
Correlation of CHEP amplitudes and pain ratings. **(A)** (left) Group (N=16) mean ± SEM N1-P1 peak-to-peak amplitudes (blue) and pain ratings (black) across the 40 trials for the Sham condition. (right) Group (N=16) scatter plot depicting the correlation between the N1-P1 peak-to-peak CHEP amplitude and pain ratings across trials for the Sham condition. r = correlation coefficient, p = p-value, black line = least-squares line. **(B)** (left) Group (N=16) mean ± SEM N1-P1 peak-to-peak amplitudes (blue) and pain ratings (black) across the 40 trials for the low-intensity focused ultrasound (LIFU) to the dorsal anterior cingulate (dACC) condition. (right) Group (N=16) scatter plot depicting the correlation between the N1-P1 CHEP amplitude and pain ratings across trials for the LIFU to dACC condition. r = correlation coefficient, p = p-value, black line = least-squares line. **(C)** (left) Group (N=16) mean ± SEM N1 amplitude (blue) and pain ratings (black) across the 40 trials for the Sham condition. (right) Group (N=16) scatter plot depicting the correlation between the N1 amplitude and pain ratings across trials for the Sham condition. r = correlation coefficient, p = p-value, black line = least-squares line. **(D)** (left) Group (N=16) mean ± SEM N1 amplitude (blue) and pain ratings (black) across the 40 trials for the LIFU to dACC condition. (right) Group (N=16) scatter plot depicting the correlation between the N1 amplitude and pain ratings across trials for the LIFU to dACC condition. r = correlation coefficient, p = p-value, black line = least-squares line.

**Figure S3.**
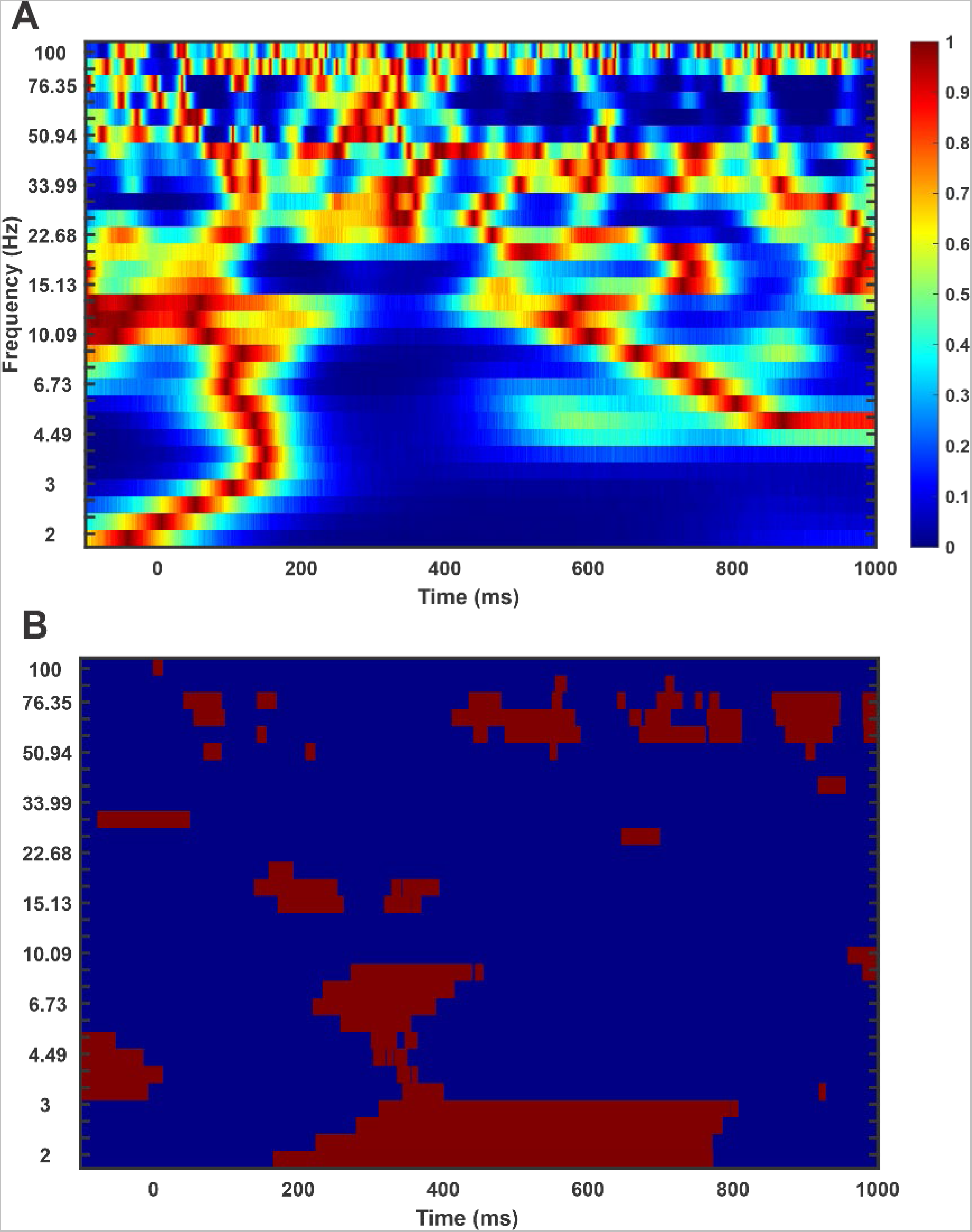
CHEP EEG time-frequency statistical maps. **(A)** Map of the raw p-values for the group (N=16) permutation testing (N=1000; p < 0.05) where warm colors represent larger p-values and cool colors represent smaller p-values. **(B)** Results from the above p-value map cluster thresholded at 10 consecutive timepoints (1 millisecond per timepoint) where red represents significant results at a cluster threshold of p < 0.05. Companion to Figure 7.

**Figure S4.**
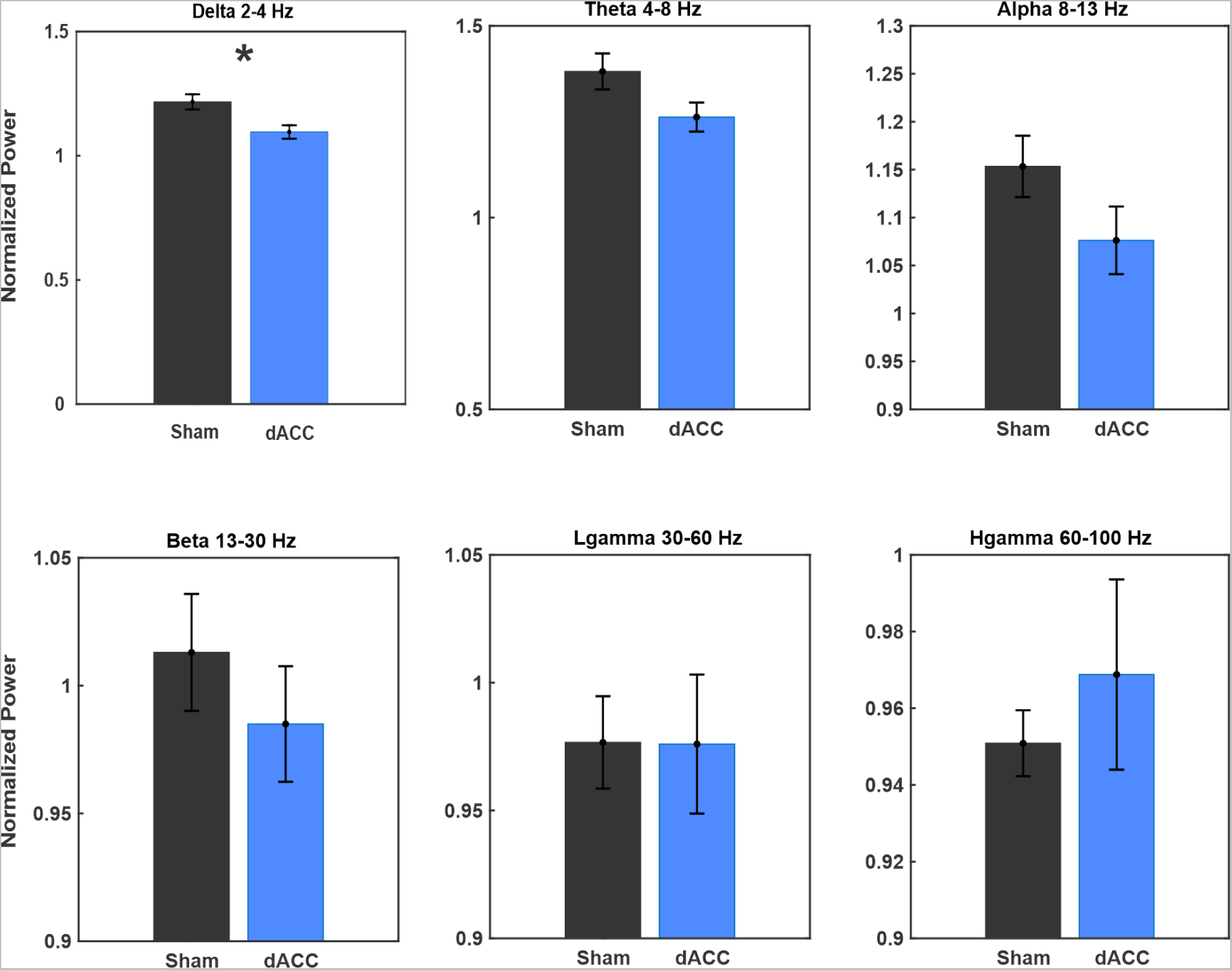
CHEP EEG frequency band power. Group (N=16) mean ± SEM normalized power for each electroencephalogram (EEG) frequency band of interest. Data was taken from +200 to +600 milliseconds (ms) from the onset of the contact heat-evoked potential (CHEP) stimulus. Frequency bands in hertz (Hz) include delta (2-4 Hz), theta (4-8 Hz), alpha (8-13 Hz), beta (13-30 Hz), low gamma (30-60 Hz) and high gamma (60-100 Hz). Black is Sham and blue is low-intensity focused ultrasound (LIFU) to the dorsal anterior cingulate (dACC) conditions. Power is represented as normalized to the baseline time window −1500ms to −500ms prior to the onset of the CHEP stimulus. Asterisk signifies significant results at p < 0.05.

**Figure S5.**
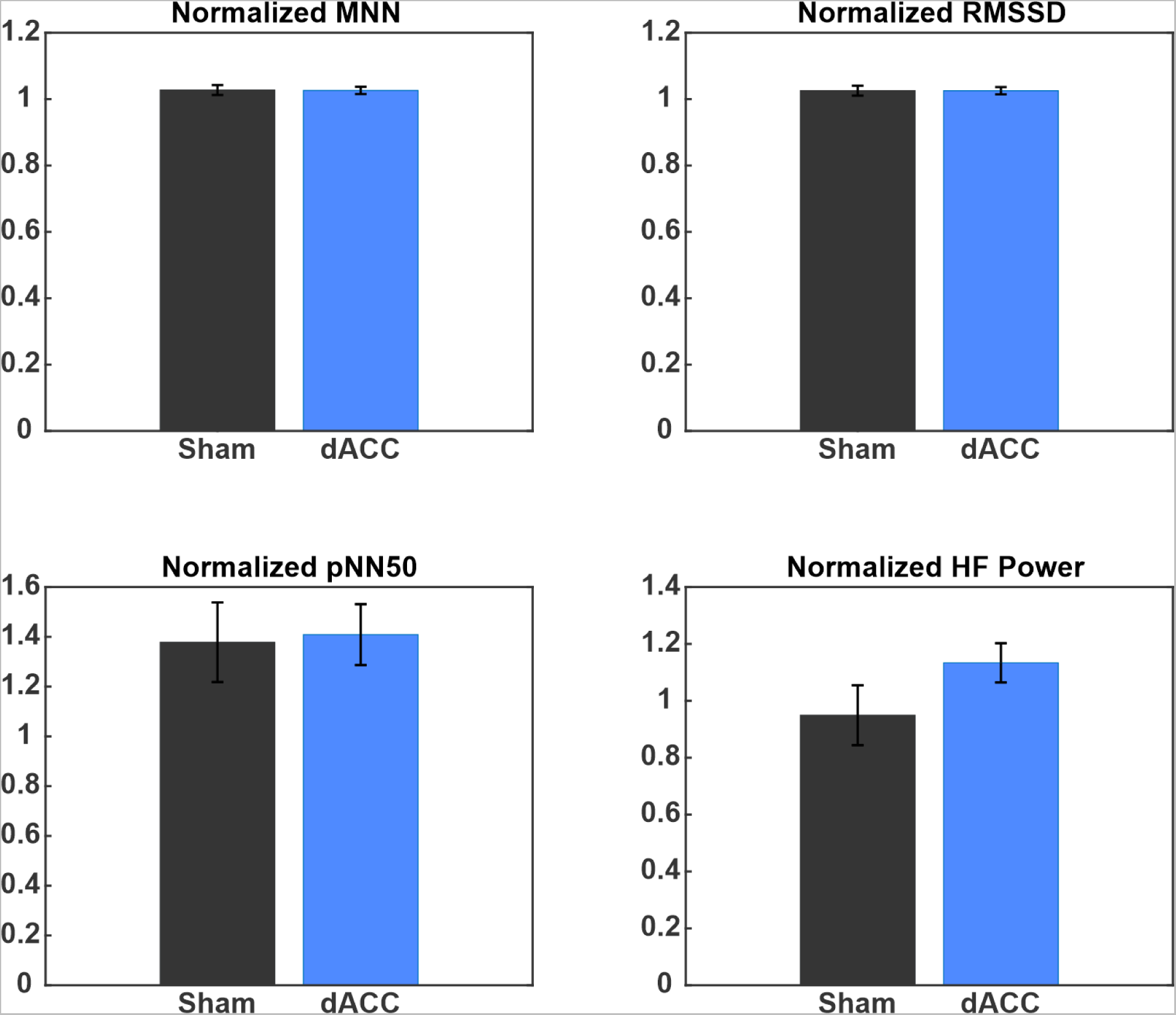
Other HRV metrics during CHEP testing. Group (N=16) mean ± SEM heart rate variability (HRV) metrics including the mean normal sinus difference (MNN) (top left), root mean square of successive differences (RMSSD) (top right), proportion of normal sinus differences greater than 50ms (pNN50) (bottom left) and high-frequency (HF) power (bottom right). Black is Sham and blue is low-intensity focused ultrasound (LIFU) to the dorsal anterior cingulate (dACC) conditions. Y-axes are the respective metrics during contact heat-evoked potential (CHEP) testing normalized to values during the 5-minute baseline period. None met statistical significance. Companion to Figure 5.

**Figure S6.**
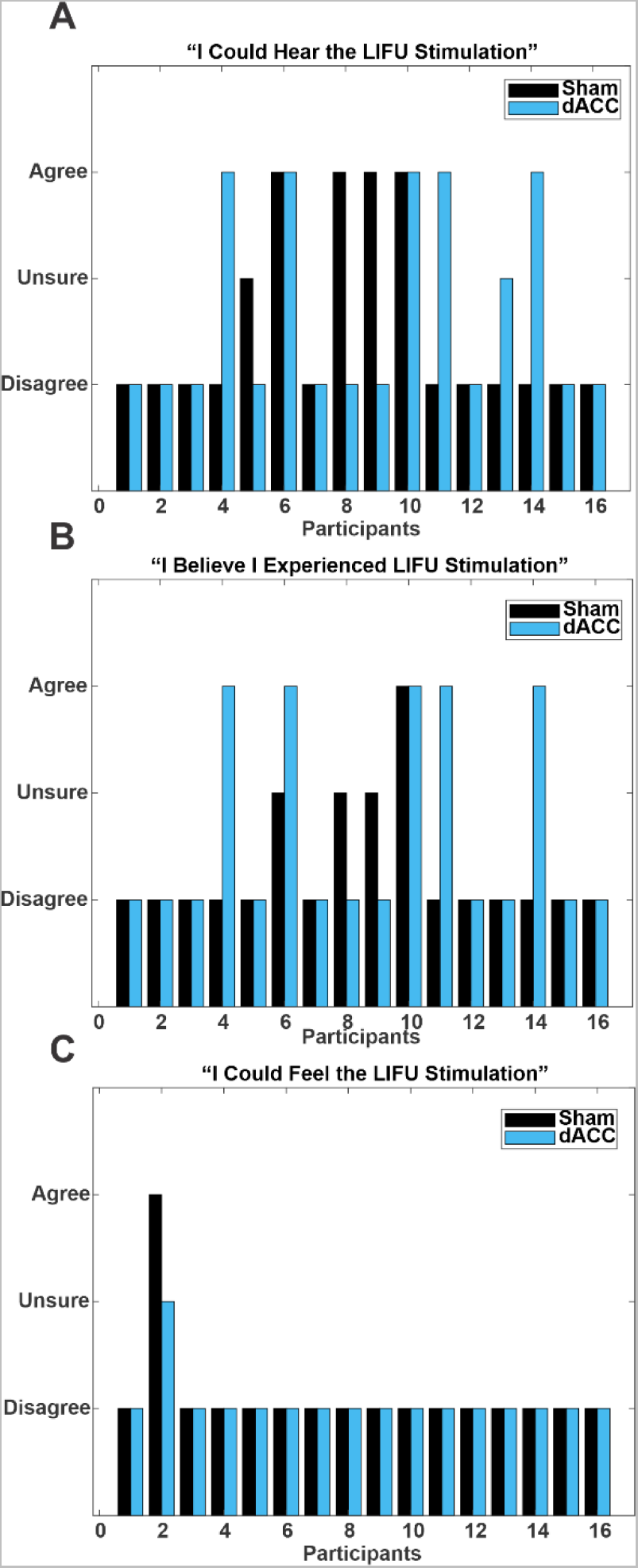
Individual subject auditory query results. **(A)** Individual subject results between Sham (black) and low-intensity focused ultrasound (LIFU) to the dorsal anterior cingulate (dACC) (blue) conditions for the question “I could hear the LIFU stimulation.” **(B)** Individual subject results between Sham (black) and LIFU to dACC (blue) conditions for the question “I believe I experienced LIFU stimulation.” **(C)** Individual subject results between Sham (black) and LIFU to dACC (blue) conditions for the question “I could feel the LIFU stimulation.” The participants on the x-axis are the same across all three questions.

**Figure S7.**
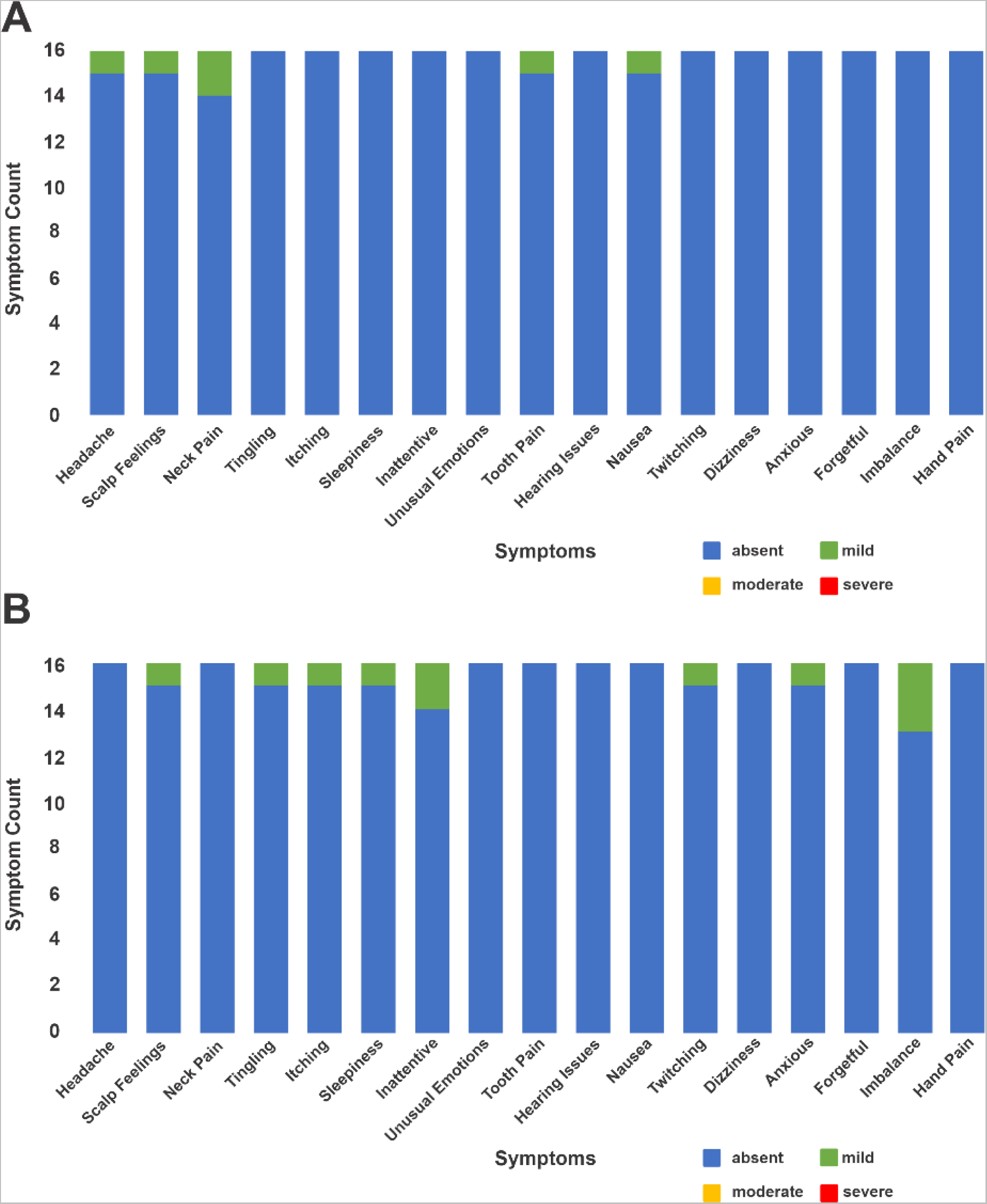
Report of symptoms questionnaire results. Report of symptoms (ROS) was administered before and after contact heat-evoked potential (CHEP) testing for both conditions. **(A)** Presence of symptoms during the low-intensity focused ultrasound (LIFU) to the dorsal anterior cingulate (dACC) condition. Each bar represents in total count for the presence of each symptom by severity across all participants (N=16). The presence of symptoms for each participant after LIFU were adjusted for the presence of symptoms prior to LIFU. No difference or fewer symptoms are represented as absent. **(B)** Presence of symptoms during the sham condition. Each bar represents in total count for the presence of each symptom by severity across all participants (N=16). The presence of symptoms for each participant after CHEP testing were adjusted for the presence of symptoms prior to CHEP testing. No difference or fewer symptoms are represented as absent.

## TABLES

**Table S1.** Individual subject depth from the scalp to the dACC target (MNI: 0,18,30) and estimated intracranial pressure in kilopascals (kPa)

| Sex | Subject | Depth to dACC Target (mm) | Estimated Intracranial Pressure (kPa) |
| --- | --- | --- | --- |
| Male | 1 | 49.3 | 53.99 |
|  | 2 | 49.2 | 52.27 |
|  | 3 | 53.7 | 37.36 |
|  | 4 | 54.5* | 48.81 |
|  | 5 | 45.6 | 47.22 |
|  | 6 | 47.8 | 77.52 |
|  | Mean (SD) | 50.02 (3.44) | 52.86 (13.40) |
| Female | 1 | 51.6 | 85.54 |
|  | 2 | 46 | 86.19 |
|  | 3 | 47 | 61.46 |
|  | 4 | 43.8 | 75.40 |
|  | 5 | 46.7 | 53.39 |
|  | 6 | 45.7 | 47.87 |
|  | 7 | 45.7 | 69.19 |
|  | 8 | 42.7 | 39.53 |
|  | 9 | 41.5 | 63.45 |
|  | 10 | 46.6 | 62.91 |
|  | Mean (SD) | 45.73 (2.76) | 64.49 (15.24) |
| Both | Mean (SD) | 47.39 (3.62) | 60.13 (15.27) |

\*secondary site (MNI:0,26,26)

